# Neutrophils repurpose the nucleolus as a cytokine reservoir and secretory organelle

**DOI:** 10.64898/2026.05.18.726057

**Authors:** Shuying Xu, Asya Smirnov, Rachel L. Kinsella, Ananda Rankin, Ashley D. Wise-Mitchell, Jessica M. Alexander, Benjamin Patty, Sophie M. Mikhail, Rachel M. Bello, Darren Kraemalmeyer, Sebastian Boluarte, Aamir Khan, Benjamin L. Allsup, Sheikh Mahmud, Bryan D. Bryson, Joshua T. Mattila, Eric L. Greer, Siyuan Ding, Regina A. Clemens, Andrew J. Monteith, Eliezer Calo, Christina L. Stallings

**Author notes:** Department of Medicine, Division of Infectious Diseases and Emory Vaccine Center, Emory University School of Medicine, Atlanta, GA, USA.

## Abstract

Type I interferons (IFN-I) are critical for antiviral defense but can drive severe pathology when dysregulated. Excess IFN-I is associated with prominent neutrophil accumulation; however, the contribution of neutrophils to IFN-I overproduction remains underexplored. In all cell types previously studied, IFN-I is synthesized de novo following sensing of microbial or host-derived inflammatory stimuli. Contrary to this paradigm, we find that neutrophils express IFNα during development and store it in the nucleolus, a membrane-less intranuclear condensate classically functioning in ribosome biogenesis. TLR-mediated bacterial sensing induces a nucleolar stress response in neutrophils that triggers rapid release of nucleoli-stored IFNα independent of de novo protein synthesis. These findings reveal that neutrophils have repurposed the nucleolus as a cytokine storage and secretory organelle, identify the first naturally occurring immunoregulatory function of nucleolar stress, and provide insight into the relationship between detrimental IFN-I levels and neutrophil accumulation.

## Main

Type I interferons (IFN-I) are central mediators of the mammalian antiviral defense. In all cell types previously studied, IFN-I is produced *de novo* upon sensing of a pathogen or damage associated molecular pattern, which triggers transcription of IFN-I genes and translation of IFN-I proteins for secretion^1,2^. IFN-I, which consists predominantly of IFNβ and numerous IFNα subtypes, signals through the IFN-α/β-receptor (IFNAR) to activate the JAK and STAT signaling pathway, inducing the expression of interferon-stimulated genes (ISG) that engage in antiviral and immunomodulatory functions^1,3^. All 13 human IFNα and 14 mouse IFNα proteins have similar structures, ranging in length of 161-167 amino acids, and exhibit as high as 75-99% conserved protein sequences within a given species^4,5^. Both immune cells and non-hematopoietic cells can produce IFN-I. However, differences in cell type and tissue-specific expression of the IFNα subtypes have been reported, implying cell specificity in the induction of IFN-I^6^.

In contrast to the critical role of IFN-I in antiviral defense, excessive IFN-I signaling can drive serious inflammatory pathology in a wide range of autoimmune diseases and non-viral infections^1,7-10^. A common feature of pathological IFN-I responses is the excessive accumulation and activation of neutrophils^11-16^. In these contexts, neutrophils are commonly viewed as responders to IFN-I, where IFN-I secreted from other cell types can promote neutrophil accumulation, swarming, and extracellular trap (NET) release^17-21^. One example is the association of IFNAR signaling in neutrophils to increased susceptibility to *Mycobacterium tuberculosis* (Mtb) infection^13-15^. Studies into the sources of IFN-I during Mtb infection have focused on late-stage disease within granulomas and have identified macrophages and plasmacytoid dendritic cells (pDCs) as the producers of IFN-I that signal to neutrophils^14,20^. However, in recent investigations with purified primary neutrophil cultures infected with Mtb *ex vivo*, we demonstrated that IFNAR-dependent NET release occurred in cultures where the only source of IFN-I could have been neutrophils^14,15^. These findings suggest that neutrophils themselves could be an unappreciated source of IFN-I during Mtb infection. Indeed, there is growing evidence that neutrophils are capable of generating IFN-I in response to Mtb infection of susceptible mouse models^20^, staphylococcal pore-forming toxins^21^, and nucleic acid Toll-like receptor (TLR) agonists^22,23^. Despite these observations, the molecular mechanisms of IFN-I secretion by neutrophils and the contribution of neutrophil-derived IFN-I to human disease have remained elusive.

### Neutrophils release IFN-I in a rapid de novo synthesis-independent manner

To investigate neutrophils as a source of IFN-I, we used an Mtb *ex vivo* infection model. Following Mtb infection of purified primary murine neutrophils *ex vivo*, peptidyl arginine deiminase 4 (PAD4) citrullinates histones to decondense the chromatin and IFNAR signaling promotes the release of this decondensed chromatin as NETs while maintaining neutrophil viability^14,15^. Similar to our observations with mouse neutrophils, when we infected primary human neutrophils purified from the peripheral blood of healthy donors with wild-type (WT) Mtb Erdman constitutively expressing GFP (Mtb-GFP)^9,14^ *ex vivo*, NET release was observed at 120 minutes post infection (mpi) using immunofluorescence confocal microscopy with antibodies specific for citrullinated histone 3 (H3Cit) (Figure 1A, top panels). Histone citrullination during Mtb infection of human neutrophils was dependent on PAD4 and inhibited by the pan-PAD inhibitor Cl-amidine (Figure 1A, bottom left panel). Treatment of human neutrophils infected with Mtb *ex vivo* with an antibody that binds IFNAR to block IFN-I signaling resulted in the accumulation of H3Cit^+^ vesicles in the cytoplasm (Figure 1A, inset) due to the inhibition of the exocytosis of chromatin-containing vesicles, and decreased extracellular NET released (Figure 1A, bottom right panel). Thus, in cultures of purified mouse^14^ or human neutrophils, IFNAR signaling is required for NET release, implying the presence of IFN-I in the neutrophil cultures. We also directly analyzed the canonical output of IFNAR signaling by monitoring the expression of ISGs in purified neutrophil cultures during Mtb infection. We isolated bone marrow neutrophils from WT C57Bl/6J mice or mice deleted for the gene encoding a subunit of IFNAR (*Ifnar1*) and performed RNA sequencing (RNA-seq) at 240 mpi with Mtb. Differential gene expression analysis revealed that many established ISGs^24^, including *Irf9, Oasl2*, and *Irgm2*, were expressed in purified neutrophil cultures during Mtb infection in an IFNAR-dependent manner (Figure 1B, Supplemental Table 1). Together, these data indicate that purified mouse and human neutrophil cultures release functional IFN-I during Mtb infection.

**Figure 1.**
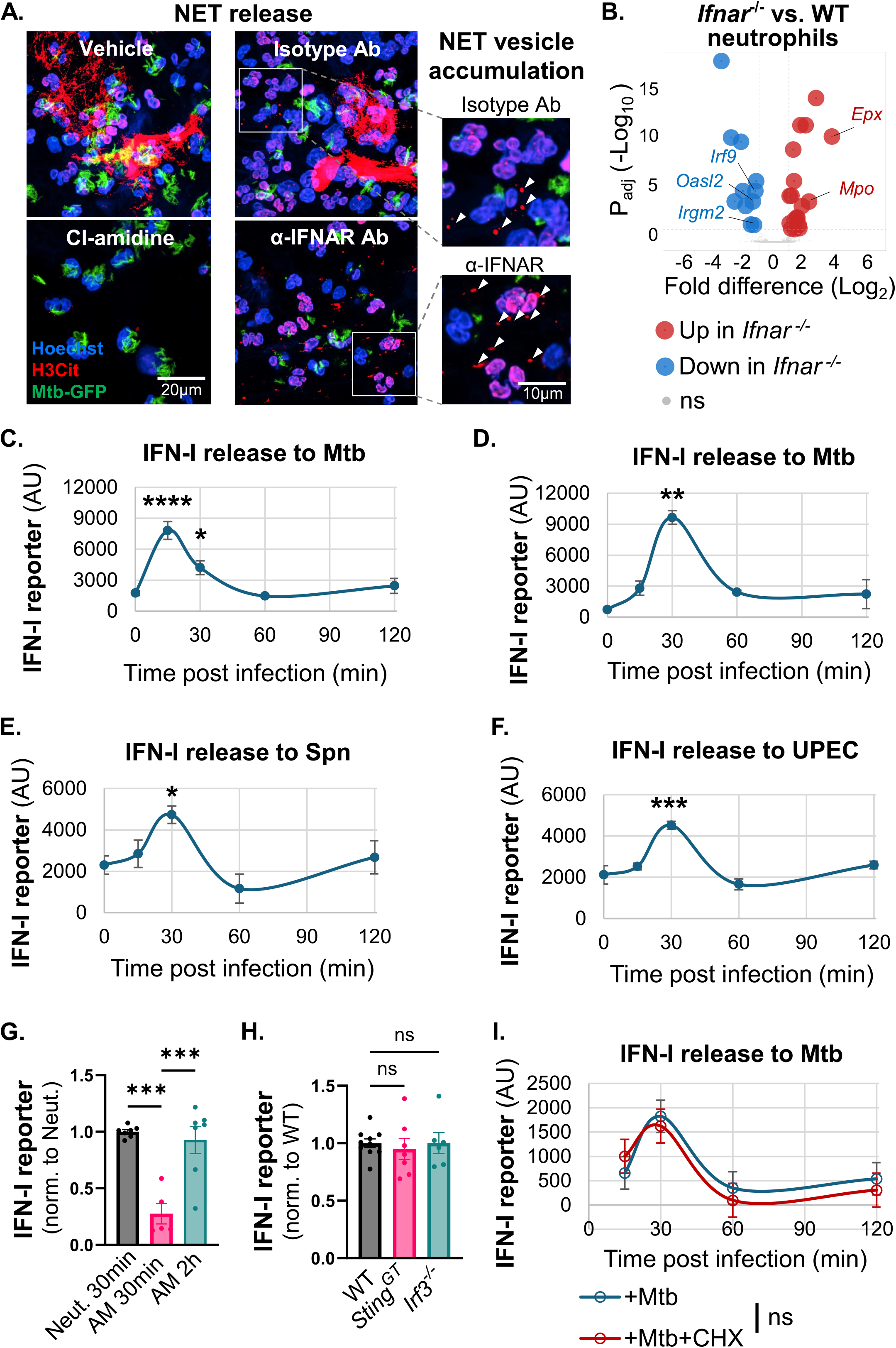
Neutrophils exposed to bacteria release functional IFN-I in a rapid *de novo* synthesis-independent manner. **A**. Immunofluorescence confocal microscopy of human neutrophils treated with vehicle (DMSO), 50 µM Cl-amidine (pan-PAD inhibitor), α-IFNAR blocking antibody, or IgG isotype control, during infection with Mtb at a multiplicity of infection (MOI) of 5. Images are from 120 mpi and show DNA (Hoechst), citrullinated histone 3 (H3Cit), and Mtb-GFP. Images on the right are zoomed in from the region in the white box. Arrowheads indicate vesicles containing citrullinated histones. **B**. Volcano plot summarizing differential gene expression from bulk RNAseq analysis of WT and *Ifnar*^*-/-*^ mouse neutrophils at 240 mpi with Mtb. Genes significantly down-(blue) or up-regulated (red) in *Ifnar*^*-/-*^ relative to WT (P_adj_ < 0.05) are shown. **C-F**. Total IFNAR-dependent luciferase activity from interferon-stimulated response element (ISRE) HeLa reporter cells exposed to supernatants collected from Mtb-infected human neutrophils (C), Mtb-infected mouse neutrophils (D), Spn-infected mouse neutrophils (E), or UPEC-infected mouse neutrophils (F). **G**. Total IFNAR-dependent luciferase activity from ISRE reporter cells exposed to supernatants collected from Mtb-infected mouse neutrophils (Neut.) or alveolar macrophages (AM) at indicated timepoints. **H**. Total IFNAR-dependent luciferase activity from ISRE reporter cells exposed to supernatants collected from WT, *Sting*^*GT*^, or *Irf3*^*-/-*^ mouse neutrophils at 30 mpi with Mtb. **I**. Total IFNAR-dependent luciferase activity from ISRE reporter cells exposed to supernatants collected from Mtb-infected human neutrophils that were pre-treated with 10 µg/mL cycloheximide as indicated. Each data point is the average of independent wells of infected cells in a representative experiment from at least two independent experiments (B-F, I), or represents an independent well of infected cells from at least two independent experiments (G-H). Error bars represent the mean ± SEM. Statistical differences to the zero timepoint or to WT neutrophil controls were determined using ordinary one-way ANOVA (C-H) or two-way ANOVA (I) with Tukey’s post-hoc test: *p < 0.05, **p < 0.01, ***p < 0.001, **** < 0.0001.

To determine the kinetics of IFN-I secretion from neutrophils during Mtb infection, we infected primary neutrophils from the blood of healthy human donors (Figure 1C) or the bone marrow of WT mice (Figure 1D) with Mtb, and collected supernatants from infected neutrophils at 15, 30, 60, and 120 mpi, during which over 90% of neutrophils remained viable through the course of infection (Supplemental Figure 1A). The supernatants were added to a reporter HeLa cell line that expresses luciferase under the control of interferon-stimulated response elements (ISREs)^25^. Supernatants were incubated with the ISRE reporter HeLa cells in the presence or absence of α-IFNAR1 blocking antibody, which allowed us to identify the luciferase activity that was dependent on IFNAR signaling as a proxy for the amount of IFN-I in the supernatant from neutrophil cultures. We found that IFN-I levels in the supernatants from both Mtb-infected mouse and human neutrophils peaked within 30 mpi (Figure 1C-D). Heat-killed Mtb was sufficient to trigger this very rapid IFN-I secretion from mouse neutrophils (Supplemental Figure 1B), demonstrating that infection with live Mtb was not required. To determine if other bacterial infections could induce a similar rapid secretion of IFN-I, we infected WT mouse neutrophils with *Streptococcus pneumoniae* (Spn) strain TIGR4 and uropathogenic *Escherichia coli* (UPEC) strain UTI89, and observed similar peaks of IFN-I release within 30 mpi (Figure 1E-F), regardless of whether we used live or heat-killed bacteria (Supplemental Figure 1C-D). The level of luciferase activity detected from the reporter HeLa cells following addition of supernatant from Mtb-infected or heat-killed Mtb-treated neutrophils was similar in magnitude to that observed following direct addition of 10 pg recombinant IFNα or IFNβ to the reporter cells (Supplemental Figure 1E).

The kinetics of IFN-I secretion from infected neutrophils was significantly faster than observed with Mtb-infected *ex vivo* cultured murine alveolar macrophages^26^, where at 30 mpi, mouse neutrophils released 4-fold more IFN-I than alveolar macrophages, which did not reach a level of secreted IFN-I comparable to that observed from neutrophils until 120 mpi (Figure 1G). In macrophages infected with Mtb, IFN-I transcription and translation are induced following sensing of Mtb by cGAS/STING and other nucleic acid sensors, which activate IRF3-dependent transcription of IFN-I genes^2,27-29^. The kinetics of IFN-I secretion from Mtb-infected neutrophils is faster than typically required to complete the processes of bacterial sensing and *de novo* IFN-I synthesis. Indeed, we found that Mtb-infected mouse neutrophils released IFN-I independently of both STING and IRF3 (Figure 1H), indicating that IFN-I secretion from neutrophils does not rely on transcriptional activation. Consistent with this, inhibition of protein synthesis by addition of cycloheximide during Mtb infection had no effect on the magnitude or kinetics of IFN-I release by human neutrophils (Figure 1I). This is in contrast to neutrophil IFN-I production triggered by poly(I:C)^22^, which required IRF3 and *de novo* protein synthesis (Supplemental Figure 1F-G). Together, these findings demonstrate that neutrophils employ a noncanonical and *de novo* synthesis-independent mechanism of IFN-I secretion upon sensing bacteria.

### Neutrophils establish a reservoir of pre-made IFNα in the nucleolus during maturation

The rapid and *de novo* synthesis-independent secretion of IFN-I from neutrophils exposed to bacteria suggests that neutrophils harbor a pre-made reservoir of IFN-I poised for rapid release. Indeed, western blotting of whole cell lysates from naïve primary mouse bone marrow neutrophils using a pan-IFNα antibody detected the presence of IFNα in naïve neutrophils but not naïve alveolar macrophages (Figure 2A), supporting that IFNα is pre-made and stored within naïve neutrophils. To determine where the IFNα is stored in neutrophils, we examined IFNα subcellular localization by immunofluorescence confocal microscopy with the pan-IFNα antibody. Because IFNα is a secreted cytokine, we initially expected the pre-made IFNα reservoir to reside in a cytoplasmic or secretory compartment. Instead, we found that naïve primary mouse and human neutrophils expressed IFNα (Figure 2B-C, top panel) that colocalized with the nucleolar marker fibrillarin within the nucleus (Figure 2B-C, bottom panel). These data demonstrate that naïve neutrophils store pre-made IFNα in their nucleoli.

**Figure 2.**
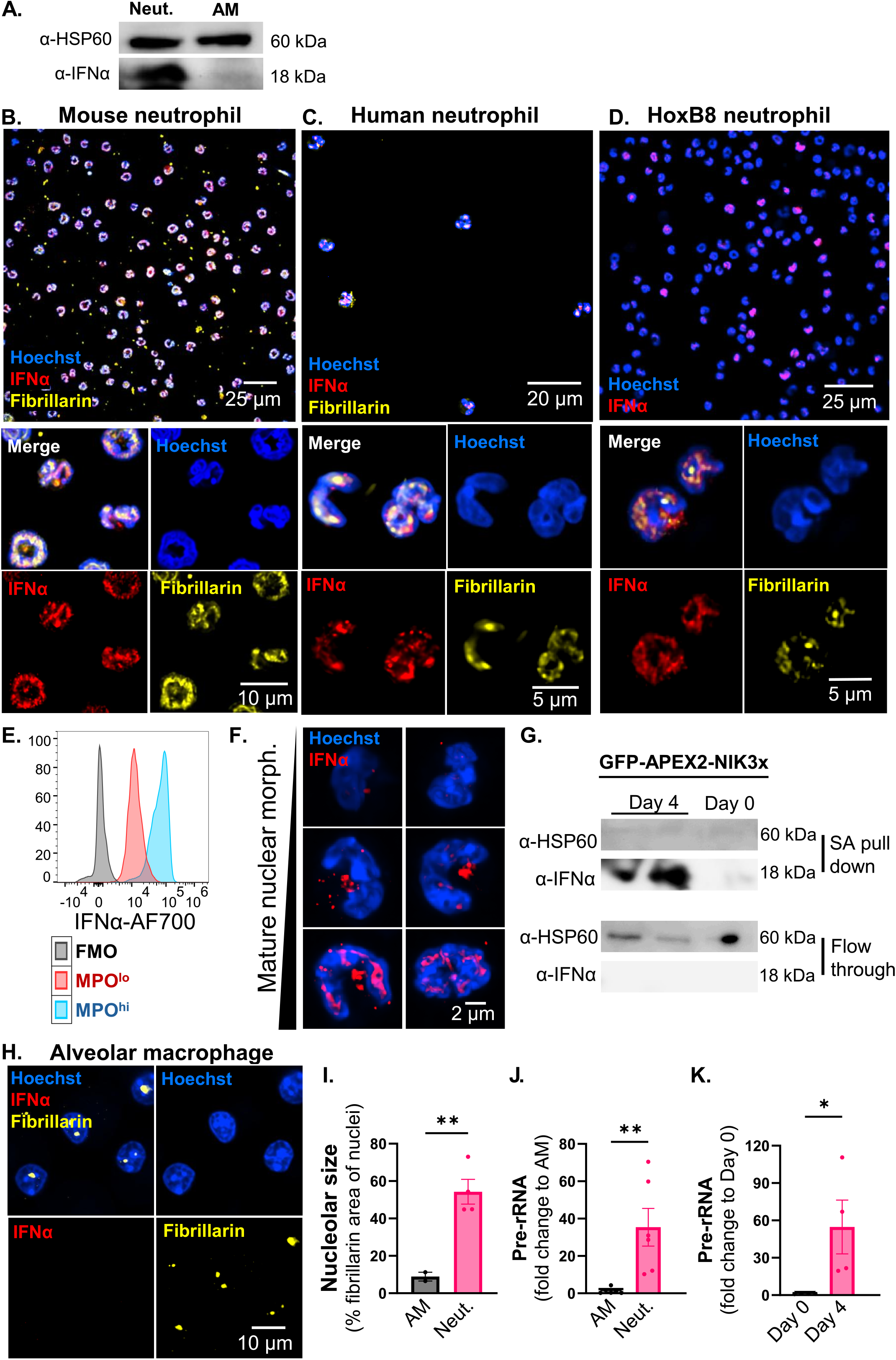
Neutrophils accumulate pre-rRNAs and establish a reservoir of pre-made IFNα in the nucleolus during maturation. **A**. Western blot for HSP60 and IFNα in whole cell lysates of naïve mouse neutrophils (neut.) or alveolar macrophages (AM). **B-D**. Immunofluorescence confocal microscopy of naïve mouse neutrophils (B), human neutrophils (C), or Hoxb8 neutrophils (D) at 20X (top panel) or 60X (bottom panel) magnification. Images show DNA (Hoechst), IFNα, and fibrillarin as indicated. **E**. FACS analysis of intracellular IFNα expression in GR-1^+^MPO^lo^ and GR-1^+^MPO^hi^ Hoxb8 neutrophils. **F**. Immunofluorescence confocal microscopy of HoxB8 neutrophils ranked by immature to mature nuclear morphology. Images show DNA (Hoechst) and IFNα. **G**. Western blot for HSP60 and IFNα in streptavidin (SA) pull down and flow through of GFP-APEX2-NIK3x-expressing HoxB8 neutrophils and progenitor cells following proximity biotinylation. **H**. Immunofluorescence confocal microscopy of mouse alveolar macrophages. Images show DNA (Hoechst), IFNα, and fibrillarin. **I**. ImageJ quantitation for percent fibrillarin area in total nuclear area of naïve mouse alveolar macrophage and neutrophils. **J-K**. qRT-PCR of 45S pre-rRNA in naïve mouse alveolar macrophage and neutrophils (J), or in HoxB8 progenitors (Day 0) and Hoxb8 neutrophils (Day 4) (K). Each data point is one field of view (I) or an independent well of infected cells (J-K) from at least two independent experiments. Error bars represent the mean ± SEM. Statistical differences between neutrophil and alveolar macrophage, or between HoxB8 progenitors and HoxB8 neutrophils, were performed using Student’s t-test: *p < 0.05, **p-value < 0.01.

To determine when neutrophils begin expressing IFNα and store it in the nucleoli, we utilized an *in vitro* system of neutrophil differentiation involving murine bone marrow GR-1^-^ neutrophil progenitors that are conditionally immortalized with ER-HoxB8 and can be induced to differentiate into mature neutrophils^30^. After 4 days of differentiation, over 80% of the HoxB8 cells express the neutrophil marker GR-1 (Supplemental Figure 2A) and 20% of the cells also express high levels of myeloperoxidase (MPO), a component of primary granules and indicative of more mature neutrophils^31^ (Supplemental Figure 2B). While undifferentiated neutrophil progenitors did not express IFNα (Supplemental Figure 2C, left panel), IFNα expression was detected by immunofluorescence microscopy and flow cytometry in 10-20% of HoxB8 neutrophils after 4 days of differentiation (Figure 2D, Supplemental Figure 2C),, and this IFNα expression was higher in the MPO^hi^ mature HoxB8 neutrophils compared to their MPO^lo^ less mature counterparts (Figure 2E). By binning the cells in the HoxB8 cultures after 4 days of differentiation based on their nuclear morphology, as a proxy of neutrophil maturity^32,33^, we found that the majority of HoxB8 neutrophils expressing IFNα harbored the IFNα in their nucleoli (Figure 2D, bottom) and exhibited highly segmented nuclear morphology indicative of more mature neutrophils (Figure 2F). In addition, we observed that less mature metamyelocytes expressed low levels of perinuclear IFNα, which we predict is what becomes sequestered into the nucleoli in band cells and more mature neutrophils (Figure 2F). Together, these data indicate that IFNα is expressed and localizes to the nucleolus as neutrophils mature. We also generated HoxB8 neutrophil progenitors expressing a GFP-APEX2-NIK3 construct that localizes to the nucleolus (Supplemental Figure 2D) and can be used to proximity label nucleolar proteins^34^. Following APEX2-mediated proximity labeling in HoxB8 neutrophil cultures before and after 4 days of differentiation, we pulled down biotinylated IFNα in the 4-day differentiated HoxB8 neutrophils but not in the undifferentiated HoxB8 neutrophil progenitor cells (Figure 2J), confirming nucleolar localization of IFNα in more mature neutrophils.

The nucleolus is a prominent nuclear condensate typically defined as the site of ribosome biogenesis. Ribosomal RNA (rRNA) synthesis and processing drive nucleolar phase separation, architecture, retention of nucleolar proteins, and function^35-40^. The nucleolar IFNα reservoir observed in neutrophils is unprecedented, and there are no other reported instances of immune mediators being stored in the nucleoli of cells. The nucleolar IFNα reservoir is also unique to neutrophils and not observed in naïve murine alveolar macrophages (Figure 2H). Therefore, we investigated whether the neutrophil nucleolus exhibited unique features that allow for IFNα storage. We noted that the neutrophil nucleoli appeared enlarged and adopted the shape of the multi-lobular nuclei of neutrophils (Figure 2B-C, “Fibrillarin”), as opposed to the round nucleoli typically described^40^ and observed in alveolar macrophages (Figure 2H, “Fibrillarin”). Based on fibrillarin staining, the nucleolus comprised over 50% of the nuclear area in neutrophils, compared to only 10% in alveolar macrophages (Figure 2I). Accumulation of unprocessed precursor rRNAs (pre-rRNAs) and the associated rRNA-binding proteins can promote nucleolar enlargement and nucleolar protein retention in other cell types^39-42^. Mature neutrophils are reported to have low levels of ribosome biosynthesis^43^, and so we inquired whether this correlated with an accumulation of unprocessed pre-rRNAs. Using quantitative real-time PCR (qRT-PCR), we detected significantly higher levels of 45S pre-rRNA in neutrophils compared to alveolar macrophages (Figure 2J). These data indicate that mature neutrophils harbor unusually large nucleoli that are associated with elevated 45S pre-rRNA accumulation and IFNα nucleolar localization.

Neutrophils acquire their multi-lobulated nuclear shape during maturation^32,44^. To examine whether the unique features of the neutrophil nucleolus also develop during neutrophil maturation, we measured 45S pre-rRNA levels in the HoxB8 neutrophil cultures before and after differentiation. 4-day differentiated HoxB8 neutrophil cultures harbored significantly higher levels of 45S pre-rRNA compared to HoxB8 neutrophil progenitors before differentiation (Fig. 2K), supporting that neutrophils accumulate pre-rRNA during maturation. To further quantify the relationship between nucleolar morphology and IFNα nucleolar localization during neutrophil maturation, we performed imaging flow cytometry of HoxB8 neutrophil cultures after 4 days of differentiation. The cell nucleolus, nucleoplasm, and cytoplasm compartments were segmented using masks generated from fibrillarin staining, Hoechst staining, and brightfield images, respectively (Supplemental Figure 2E). We quantified the percentage of IFNα signal in each subcellular compartment and found that the majority of IFNα^+^ cells exhibited over 70% of IFNα staining within the nucleolus mask, where IFNα colocalized with fibrillarin (Supplemental Figure 2F-G). In contrast, less than 10% and 30% of IFNα was detected in the nucleoplasm and cytoplasm, respectively (Supplemental Figure 2G). We applied machine learning to identify features in Hoechst or brightfield images that best differentiated between cells with high versus low levels of IFNα nucleolar localization. The top features correlating with nucleolar IFNα localization were a parameter for high nuclear segmentation (Intensity_Valley), when performing analysis on all features, and a parameter for high nuclear elongation (Length_Object), when performing an analysis restricted to size and shape features (Supplemental Figure 2H), supporting that IFNα localization to the nucleolus correlates with the elongated and lobular nuclei observed in mature neutrophils. These data suggest that during neutrophil maturation, the nucleolus accumulates pre-rRNA and enlarges, at which point the IFNα that has been expressed during maturation is sequestered within it. Thus, the unique architecture of the neutrophil nucleolus may support the organization of an IFNα storage condensate.

### TLR signaling triggers neutrophils to release nucleolar IFNα

To determine whether the nucleoli-localized IFNα in naïve mature neutrophils is the source of IFN-I released during Mtb infection, we infected mouse (Figure 3A) and human (Figure 3B) neutrophils with Mtb and imaged the infected cells at 0, 30, and 60 mpi to assess localization of IFNα. Within 30 mpi, nucleoli IFNα delocalized from the nucleus and transitioned into the cytoplasm (Figure 3A-B). By 60 mpi, IFNα was barely detectable inside the neutrophils (Figure 3A-B), which follows the kinetics of when IFN-I is detected in supernatants (Figure 1C-D). Image quantification of IFNα signal within the nucleus confirmed a significant decrease in IFNα nuclear localization by 30 and 60 mpi in both mouse (Figure 3C) and human neutrophils (Figure 3D). Imaging flow cytometry analysis confirmed that by 30 minutes post exposure to heat-killed Mtb, the majority of neutrophils harbored over 60% of the IFNα signal within the cytoplasm, with less than 40% of the IFNα signal remaining nuclear (Supplemental Figure 3A-C). Treatment of Mtb-infected human neutrophils with cycloheximide to block protein translation did not affect the disappearance of nucleolar IFNα and the appearance of IFNα in the cytoplasm (Figure 3D-E), implying that the IFNα in the cytoplasm was not synthesized *de novo* and was likely the same IFNα originally stored in the nucleolus. In contrast, when we infected murine alveolar macrophages, which do not express IFNα in the naïve state (Figure 2D; 3F, “0 min”), IFNα was not detectable until 60 mpi and this IFNα was only observed in the cytoplasm (Figure 3F). In addition, cycloheximide completely blocked the appearance of IFNα in the cytoplasm of Mtb-infected alveolar macrophages, confirming that alveolar macrophages rely on *de novo* protein synthesis following Mtb infection to express and secrete IFNα (Figure 3F, “60 min +CHX”). Our data suggest that neutrophils undergo a noncanonical IFNα secretion process that is conceptually reminiscent of degranulation, the rapid release of antimicrobial and proinflammatory proteins that are pre-packaged in neutrophil granules^31,44^. However, instead of being stored in neutrophil granules, IFNα is pre-packaged into the nucleolus.

**Figure 3.**
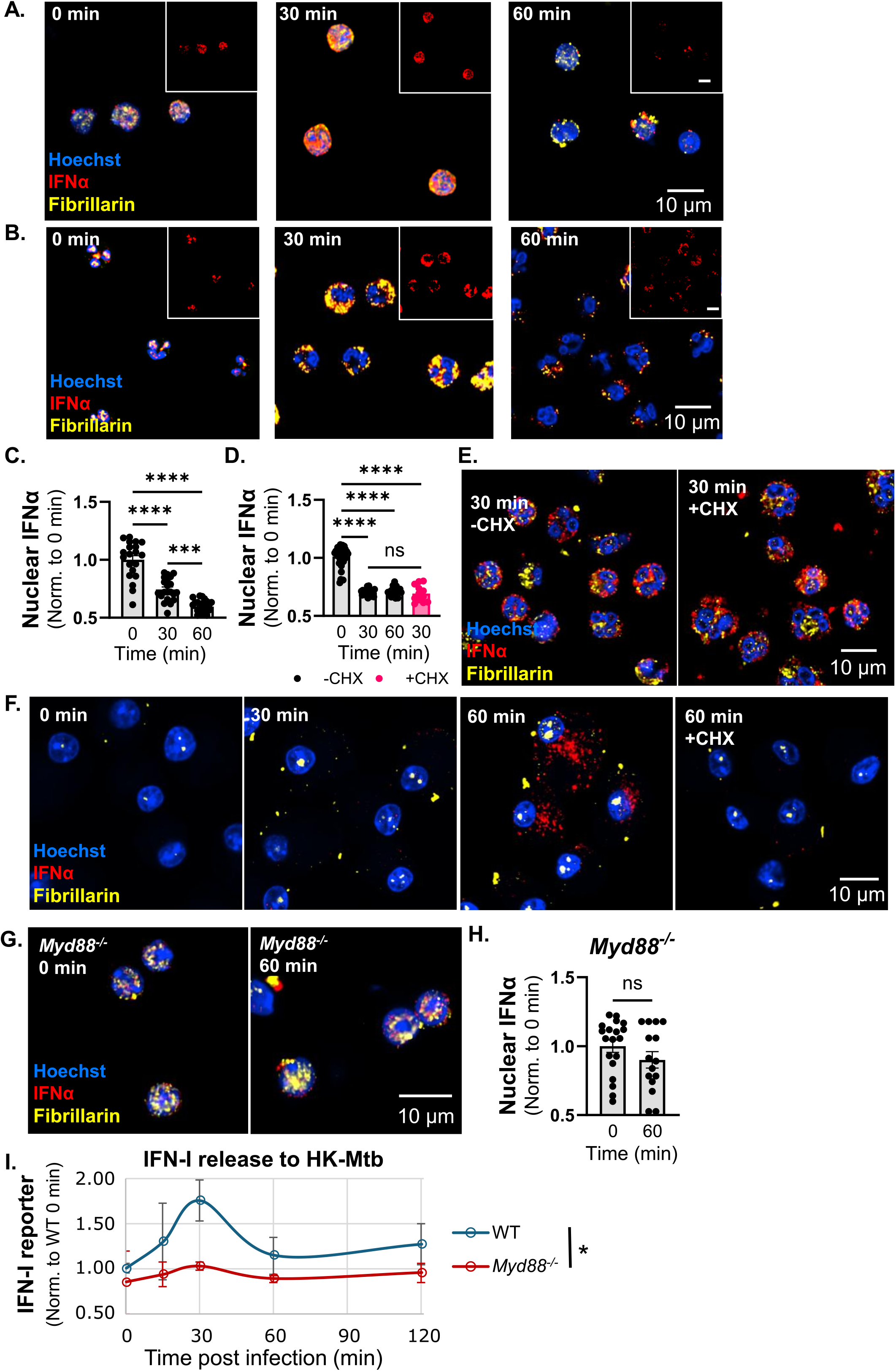
MyD88-dependent TLR signaling triggers neutrophils to release nucleoli-stored IFNα into the cytoplasm for subsequent secretion. **A-E**. Immunofluorescence confocal microscopy of mouse neutrophils (A) or human neutrophils (B, E), infected with Mtb in the absence (A, B) or presence (E) of 10 µg/mL cycloheximide. Images show DNA (Hoechst), IFNα, and fibrillarin. Insets show the IFNα channel. ImageJ quantitation of nuclear IFNα intensity in mouse (C) and human neutrophils (D). **F**. Immunofluorescence confocal microscopy of mouse alveolar macrophages infected with Mtb in presence of 10 µg/mL cycloheximide as indicated. Images show DNA (Hoechst), IFNα, and fibrillarin. **G-H**. Immunofluorescence confocal microscopy of *Myd88*^*-/-*^ mouse neutrophils treated with heat-killed Mtb. Images show DNA (Hoechst), IFNα, and fibrillarin (G). ImageJ quantitation of nuclear IFNα intensity (H). **I**. Total IFNAR-dependent luciferase activity from ISRE reporter cells exposed to supernatants collected from WT or *Myd88*^*-/-*^ mouse neutrophils treated with heat-killed Mtb. Each data point is one field of view (C-D,H) from at least two independent experiments, or the average of independent wells of infected cells from two independent experiments (I). Error bars represent the mean ± SEM. Statistical differences between time points, treatments, or genotypes were determined using Student’s t-test (H), ordinary one-way ANOVA (C-D), or two-way ANOVA (I) with Tukey’s post-hoc test: *p < 0.05, ***p < 0.001, ****p < 0.0001.

We observed the rapid release of IFN-I from neutrophils in response to live Mtb, Spn, UPEC, as well as heat-killed bacteria (Figure 1C-F, Supplemental Figure 1B-D), suggesting that the trigger for IFN-I release involves early sensing of a shared pathogen-associated molecular pattern. Among the first pattern recognition receptors on innate immune cells encountered by bacteria are TLRs, and all TLRs except TLR3 use MyD88 as a signaling adaptor to trigger immune cell activation and inflammation^45,46^. We thus investigated a potential role for MyD88 signaling in bacterial induction of neutrophil IFN-I release by exposing neutrophils isolated from *Myd88*^*-/-*^ mice to heat-killed Mtb and monitoring IFNα localization by immunofluorescence microscopy and IFN-I secretion using the ISRE HeLa reporter cells. MyD88-deficient neutrophils retained the stored IFNα in the nucleoli through 60 minutes (Figure 3G-H) and secreted significantly lower amounts of IFN-I over the 120-minute period (Figure 3I) following exposure to heat-killed Mtb compared to WT neutrophils. Thus, sensing of bacteria by neutrophils through MyD88-dependent TLRs triggers rapid redistribution of nucleolar IFNα to the cytoplasm and subsequent extracellular release.

### Nucleolar stress drives IFNα release from neutrophils

We noted that concurrent with IFNα exiting the nucleolus following Mtb infection, the nucleolar marker fibrillarin also translocated into the cytoplasm (Figure 3A-B), indicative of perturbations to the nucleolar architecture during IFNα release. Although biology of the nucleolus remains largely unexplored in neutrophils^43^, in other cells, disruption of pre-rRNA synthesis and processing results in “nucleolar stress”. Nucleolar stress, traditionally viewed as a harmful consequence of impaired ribosome production with no known physiological role, alters the architecture of nucleolar phases and redistributes nucleolar proteins within the nucleus^39,47-49^. We investigated whether IFNα release from the neutrophil nucleolus exhibited features of nucleolar stress by measuring pre-rRNA levels following exposure to heat-killed Mtb. Pre-rRNA levels decreased significantly following exposure of mouse and human neutrophils to heat-killed Mtb (Figure 4A). In contrast, exposure to heat-killed Mtb did not affect pre-rRNA levels in alveolar macrophages (Figure 4A), indicating that sensing of Mtb results in decreased pre-rRNA levels specifically in neutrophils. In comparison, treatment of neutrophils or alveolar macrophages with 50nM actinomycin D, which is known to inhibit pre-rRNA synthesis and induce nucleolar stress^50^, resulted in a significant decrease in pre-rRNA in both cell types (Figure 4B). A well-established nucleolar stress response protein that is normally localized to the nucleolus and redistributes to the nucleoplasm during nucleolar stress is the RNA helicase DDX21^47,49^. DDX21 localized to the nucleolus in naïve human neutrophils (Figure 4C) but was released into the cytoplasm along with fibrillarin and IFNα by 30 minutes of exposure to heat-killed Mtb (Figure 4D), supporting that Mtb induces disruption of the nucleolar architecture and redistribution of nucleolar proteins to the cytoplasm.

**Figure 4.**
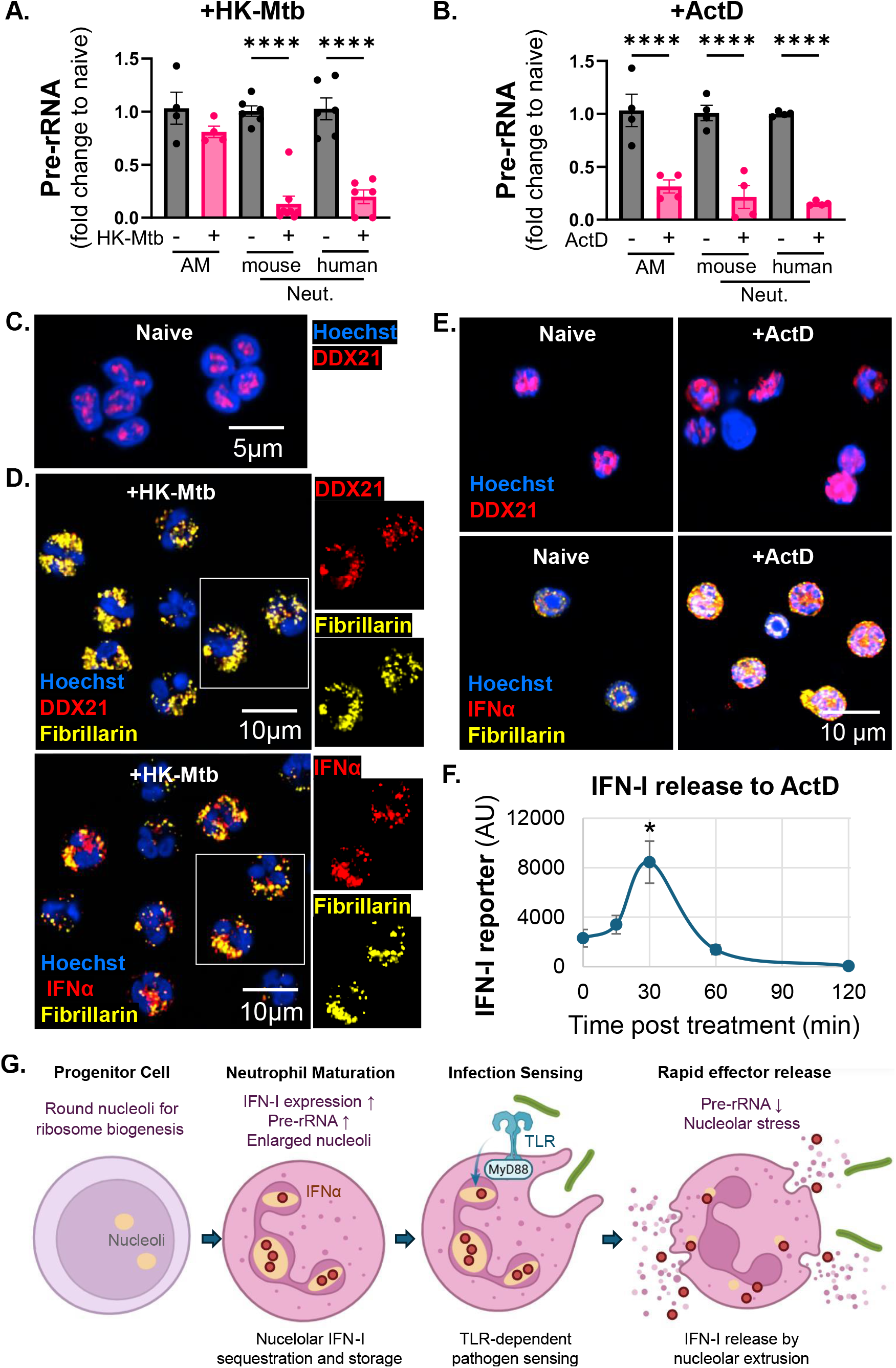
Nucleolar stress decreases pre-rRNA levels and triggers nucleolar extrusion in neutrophils to secrete IFN-I. **A-B**. qRT-PCR of 45S (mouse) or 47S (human) pre-rRNA in mouse alveolar macrophage (AM), mouse neutrophils, or human neutrophils treated with heat-killed Mtb for 30 minutes (A), or with 50 nM actinomycin D for 60 minutes (B). **C-D**. Immunofluorescence confocal microscopy of human neutrophils naïve (C) or heat-killed Mtb-treated for 30 minutes (D). Images show DNA (Hoechst), IFNα, fibrillarin, or DDX21 as indicated. **E**. Immunofluorescence confocal microscopy of mouse neutrophils naïve or treated with 50 nM actinomycin D for 30 minutes. Images show DNA (Hoechst), IFNα, fibrillarin, or DDX21 as indicated. **F**. Total IFNAR-dependent luciferase activity from ISRE reporter cells exposed to supernatants collected from mouse neutrophils treated with 50 nM actinomycin D. **G**. Model of IFN-I nucleolar storage upon neutrophil maturation and rapid release via nucleolar extrusion following infection. Each data point is an independent well of infected cells from at least two independent experiments (A-B), or the average of independent wells of infected cells in a representative experiment from at least two independent experiments (F). Error bars represent mean ± SEM. Analysis of statistical differences to naïve cells was performed using ordinary one-way ANOVA with Tukey’s post-hoc test: *p < 0.05, ****p < 0.0001.

These data indicate that bacterial sensing by TLRs, which is not a typical nucleolar stress inducer, triggers a form of nucleolar stress in neutrophils that results in a rapid decrease in pre-rRNA and redistribution of IFNα and other nucleolar proteins. Unlike classical nucleolar stress where nucleolar proteins remain confined to the nucleus, this response converts the neutrophil nucleolus into a secretory organelle, releasing nucleolar proteins into the cytoplasm prior to their subsequent extracellular release. We investigated whether a canonical inducer of nucleolar stress, actinomycin D, would also induce release of nucleolar contents into the cytoplasm of neutrophils, or if this was specific for the process initiated by sensing of bacteria. We treated mouse neutrophils with actinomycin D and monitored the effect on DDX21, fibrillarin, and IFNα subcellular localization. After 30 minutes of actinomycin D treatment, all three proteins transitioned from being localized to the nucleoli to being detected in the cytoplasm in mouse neutrophils (Figure 4E). Similarly, actinomycin D treatment for 30 minutes induced fibrillarin and IFNα to transition from the nucleolus to the cytoplasm of human neutrophils (Supplemental Figure 4A). In both mouse and human neutrophils, IFN-I was detected in supernatants by 30 minutes following treatment with actinomycin D (Figure 4F and Supplemental Figure 4B). Therefore, neutrophils not only respond to bacterial stimuli as a nucleolar stress inducer, but also uniquely respond to nucleolar stress by redistributing nucleoli-localized proteins to the cytoplasm, where, in the case of IFN-I, they can be secreted. Together, these data support a model (Figure 4G) where sensing of bacteria through TLRs induces a specialized form of nucleolar stress in neutrophils, which drives a process we are terming nucleolar extrusion, resulting in rapid release of IFN-I and other nucleolar contents.

## Discussion

Neutrophils are the first responders of the innate immune system, arriving immediately at a site of damage or infection to produce a rapid response. There are multiple unique structural and functional features of neutrophils that facilitate the speed of response. The multilobed nucleus allows neutrophils to transmigrate through endothelial and epithelial barriers to extravasate into tissues faster than other immune cells^38^. Neutrophils also synthesize granules pre-packaged with antimicrobial and proinflammatory proteins during maturation to allow for rapid release through degranulation in response to infections^31^. We report a previously unrecognized aspect of neutrophil biology that facilitates this immediate inflammatory response, by pre-rRNA- and nucleolar stress-orchestrated storage and release of effectors in the nucleolus. The mechanisms of this unique nucleolar storage and release pathway in neutrophils remain to be carefully explored. For instance, unlike canonical nucleolar stress that results in nucleolar proteins redistributing within the nucleus^47,49^, neutrophil nucleolar stress allows nucleolar components to extrude into the cytoplasm and permits subsequent extracellular release. Nucleolar extrusion in neutrophils connects TLR signaling with nucleolar stress to convert the neutrophil nucleolus into an infection-responsive secretory organelle. We propose that this nucleolar extrusion is possible because mature neutrophils have a shorter half-life than ribosomes^44,51^, allowing them to retain sufficient translational capacity while using the nucleolus and the nucleolar stress response as an innate immune effector organelle and pathway.

Our studies highlight IFN-I in particular as being stored in the neutrophil nucleolus for secretion via nucleolar extrusion, identifying neutrophils as a significant source of IFN-I. Elevated IFN-I is often associated with neutrophil-dominated inflammation in a wide range of infectious and autoimmune diseases^15,52,53^. However, neutrophils are often not recognized as a source of the IFN-I, as their rapid release of IFN-I could be missed when analyzing later stages of infection. As neutrophils are among the first cells recruited to sites of infection and damage, the rapid nature of neutrophil IFN-I release through nucleolar extrusion could condition the environment early on and affect the responses of other immune and non-immune cells at the site. In addition, recently matured neutrophils can be continually recruited from the bone marrow in large numbers, which would contribute to sustained IFN-I levels during chronic inflammatory disease^1,7,54^. IFN-I promotes other inflammatory effector functions of neutrophils, such as the release of NETs^14,18,23^, which will then be sensed by nucleic acid sensors in surrounding cells, such as macrophages and pDCs, to induce further IFN-I production from these cells^19,20^ and generate a feedforward loop of inflammation. The mechanism of nucleolar extrusion reveals how neutrophils rapidly spark the IFN-I inflammatory loop and opens new avenues for developing methods to target this inflammatory process.

## Acknowledgments

We thank Dr. Jeffery P. Henderson, Dr. Scott J. Hultgren, Dr. L David Sibley, Dr. Michael Diamond, Dr. Ashley Steed, Dr. Robert D. Schreiber (Washington University in St. Louis School of Medicine, MO), Dr. John M. Leong, Dr. Andrew Camilli, Dr. Shumin Tan (Tufts University School of Medicine, MA), Dr. David Sykes (Massachusetts General Hospital, MA), Dr. Katherine A. Fitzgerald (UMass Chan Medical School, MA) and Dr. Tadasugu Taniguchi (University of Tokyo, Japan) for generously sharing bacterial strains, cell lines, mice, reagents, and guidance. We acknowledge the assistance of Dennis Oakley and Dr. De Chen at the Washington University Center for Cellular Imaging (WUCCI), and of Wandy Beatty at the Molecular Microbiology Imaging Facility, for microscopy studies. We thank the support for spectral flow cytometry and imaging flow cytometry studies from the Mycobacteria-focused Program for Research & Innovation in Science and Medicine (MycoPRISM), Center for Women’s Infectious Disease Research (cWIDR), and Department of Molecular Microbiology Flow Cytometry Facility at Washington University School of Medicine. We also acknowledge the Genome Technology Access Center (GTAC) at the McDonnell Genome Institute (MGI), supported by the Alvin J. Siteman Cancer Center at Washington University School of Medicine and Barnes-Jewish Hospital in St. Louis, MO, and the Institute of Clinical and Translational Sciences (ICTS) at Washington University in St. Louis, for providing RNA sequencing and analysis service. Model figures are created with BioRender.com.

## Fundings

This research is supported by

National Institutes of Health grant R01 AI179024 (C.L.S.)

National Institutes of Health grant R01 AI184666 (C.L.S., B.D.B., J.T.M.)

Burroughs Wellcome Fund Investigators in the Pathogenesis of Infectious Disease Award (C.L.S.)

Potts Memorial Foundation postdoctoral fellowship (S.X.)

WUCCI is supported by the Washington University School of Medicine and The Children’s Discovery Institute of Washington University and St Louis Children’s Hospital (CDI-CORE-2015-505 and CDI-CORE-2019-813). Images were generated on a Nikon AX-R Confocal Microscope purchased with support from the Office of Research Infrastructure Programs (ORIP), a part of the NIH Office of the Director under grant OD030233.

The Siteman Cancer Center is supported in part by an NCI Cancer Center Support Grant #P30 CA091842, and the ICTS is funded by the National Institutes of Health’s NCATS Clinical and Translational Science Award (CTSA) program grant UL1 TR002345.

## Author contributions

C.L.S., E.C., and S.X. conceived and designed this study. S.X. performed experiments, analyzed data, and drafted the manuscript. C.L.S., E.C., B.D.B, J.T.M., and S.X. acquired funding support. A.S., R.L.K., A.R., A.D.W., S.B., A.J.M., and R.A.C optimized neutrophil isolation and infection experiments. A.R. performed the RNA sequencing, and A.R., B.P., S.M.M., and R.M.B. performed sequencing analysis. R.L.K., J.M.A., D.K., and S.D. performed and supported experiments using various mouse genotypes. B.L.A., S.M., A.K., E.G., and B.D.B. generated plasmids and optimized protocols used in nucleoli proximity labeling and western blotting experiments. C.L.S., A.S., A.J.M., B.L.A., S.D., A.M.J., and E.C. edited the manuscript.

## Competing interests

The authors declare that there are no competing interests.

## Data availability

The raw RNA sequencing data have been deposited in the Gene Expression Omnibus under accession number GSE330411. All other data will be deposited to Zenodo and be available from the corresponding authors by request.

## Materials and Methods

### Bacterial strains and culturing

#### *Mycobacterium tuberculosis* (Mtb)

Erdman expressing GFP (GFP-Mtb)^9,14^ was used in all experiments except for the bulk RNA-sequencing experiment and generation of heat-killed Mtb where WT Erdman was used. Mtb was cultured at 37□°C in 7H9 (Difco) medium supplemented with 10% oleic acid/albumin/dextrose/catalase (OADC), 0.5% glycerol, and 0.05% Tween 80. For use in infection of cells, GFP-Mtb grown to log-phase growth (OD600□=□0.5–0.8), in the presence of 20□µg/ml kanamycin to ensure plasmid retention, was washed with PBS, sonicated to disperse clumps, pelleted at 55□×□g for 10□minutes to remove clumps, and diluted in phosphate buffered saline (PBS) to appropriate concentrations as required. Heat-killed Mtb Erdman was prepared by growing the culture to log-phase growth before boiling at 95°C for 30-45 minutes and autoclaving. Heat-killed cultures were lyophilized for 24-48 hours, stored at 4 °C, and resuspended at a concentration of 250 µg/ml in PBS for use.

#### Serotype 4 *Streptococcus pneumoniae* (Spn)

TIGR4^55,56^ was a gift from Dr. John M. Leong and Dr. Andrew Camilli (Tufts University School of Medicine, MA). Mid-exponential growth phase aliquots of Spn were grown in Todd-Hewitt broth (BD Biosciences) supplemented with 0.5% yeast extract and 5 µl/ml Oxyrase (Oxyrase), at 37□°C in 5% CO_2_, and frozen in growth media with 20% (v/v) glycerol. Bacterial titers in aliquots were confirmed by plating serial dilutions on Tryptic Soy Agar plates supplemented with 5% sheep blood (Northeast Laboratory Services). For use in infection of cells, frozen aliquots were thawed, washed, and diluted in PBS to appropriate concentrations as required. Heat-killed Spn was prepared by growing a culture to log-phase growth before boiling at 95°C for 30-45 minutes.

#### Uropathogenic *Escherichia coli* (UPEC)

strain UTI89^57,58^ was a gift from Dr. Jeffery P. Henderson and Dr. Scott J. Hultgren (Washington University in St. Louis School of Medicine, MO). Aliquots of UPEC were grown in lysogeny broth (BD Biosciences) overnight at 37°C, measured by OD600 to determine concentration, and frozen in growth media with 20% (v/v) glycerol. For use in infection of cells, frozen aliquots were thawed, washed, and diluted in PBS to appropriate concentrations as required. Heat-killed UPEC was prepared by growing a culture overnight before boiling at 95°C for 30-45 minutes.

### Mice

Adult mice (age 6-15 weeks) of both sexes were used and randomized. C57BL/6J (000664) and *Sting*^GT^ (017537) mice were purchased from Jackson Laboratory (Bar Harbor, ME). The C57BL/6J congenic, backcrossed, and rederived *Ifnar1*^*-/-*^ mice^58,59^ were obtained from Dr. Ashley Steed (Washington University in St. Louis School of Medicine, MO), and *Myd88*^*-/-*^ mice^60,61^ from Dr. Robert D. Schreiber (Washington University in St. Louis School of Medicine, MO). *Irf3*^*-/-*^ mice^62^ were kindly provided by Dr. Katherine A. Fitzgerald (UMass Chan Medical School, MA) and Dr. Tadasugu Taniguchi (University of Tokyo, Japan). All mouse colonies are maintained in a specific pathogen-free barrier facility at Washington University in St. Louis. All procedures involving animals were conducted following the National Institute of Health guidelines for housing and care of laboratory animals and performed in accordance with institutional regulations after protocol review and approval by the Institutional Animal Care and Use Committee of Washington University in St. Louis School of Medicine.

### Infection of neutrophils or alveolar macrophages

Primary mouse neutrophils were harvested from mouse bone marrows using the EasySep Mouse Neutrophil Enrichment Kit (Stemcell Technologies) as per the manufacturer’s instructions, yielding >90% purity with less than 10% residual monocytes and lymphocytes. Whole blood obtained from healthy human volunteers under an IRB-approved protocol was used to isolate human peripheral blood neutrophils using the EasySep Direct Human Neutrophil Isolation kit (Stemcell Technologies), yielding >95% purity. Neutrophils were isolated into Hanks’ Balanced Salt Solution (HBSS) with 1mM EDTA, and counted with Trypan exclusion using a Countess 3 automated cell counter (Invitrogen). 1×10^6^ neutrophils were transferred to each well of a 24-well plate containing 12□mm glass coverslips (Fisher Scientific, #1 thickness) for infection.

Alveolar macrophages were collected from C57BL/6 mice by bronchoalveolar lavage, plated in 10 cm tissue culture-coated round dishes, and cultured in RPMI 1640 media supplemented with 10% fetal bovine serum (FBS), 1% penicillin/streptomycin, TGFβ1 (10 ng/mL, Peprotech), GM-CSF (30 ng/mL, Peprotec), and rosiglitazone (1 μM, Peprotech)^26^. Cells are maintained in 5% CO_2_ at 37°C, and at confluency, were washed with PBS and lifted with 20mM lidocaine in PBS to be passaged or used in experiments. One day prior to infection, 1×10^6^ alveolar macrophages were seeded onto 12□mm glass coverslips in a 24-well plate and allowed to attach.

To infect cells with bacteria, Mtb at a multiplicity of infection (MOI) of 5, Spn at an MOI of 2, and UPEC at an MOI of 5 were added to neutrophils or alveolar macrophages in the 24-well plates. To treat cells with heat-killed bacteria, 40 µl of heat-killed Mtb, Spn, or UPEC suspension was added per well. After incubating in 5% CO_2_ at 37□°C for the indicated time points, supernatants were collected for use in ISRE reporter cell assays, and coverslips were placed in 4% PFA fixation for immunofluorescence (IF) microscopy analysis. For pan-PAD inhibitor treatment, 50 µM Cl-amidine (Sigma) was added to each well of infected cells. For IFNAR1- blocking antibody (Leinco Technologies, Clone MAR1-5A3, I-401) or normal mouse IgG isotype control (Leinco Technologies, N229) treatment, 10 µg/ml of respective antibody was added to each well of infected cells. For actinomycin D (Sigma) treatment, 50 nM actinomycin D was added to each well of cells. For microscopy readout of cycloheximide (Goldbio) treatment, 10 µg/ml of cycloheximide was added directly to each well of infected cells at the start of the infection. For the interferon-stimulated response element (ISRE) reporter assay readout of cycloheximide treatment, 10 µg/ml of cycloheximide was pre-incubated with cells for 30 minutes and removed before the start of infection. For poly(I:C) stimulation of *de novo* IFN-I synthesis in neutrophils^22^, 0.25 µg of poly(I:C) in 100 µl of OptiMem and 2 µl of Lipofectamine 2000 (ThermoFisher) was added to each well of cells.

### Interferon-stimulated response element (ISRE) reporter cell assay

HeLa cells stably expressing 11×-ISRE-Gaussia Luciferase were generated and maintained as described previously^25^. Briefly, cells were cultured in RPMI1640 with 10% FBS in 5% CO_2_ at 37 °C. To perform IFN-I-activated luciferase reporting, the ISRE reporter cells were plated in 96-well plates at 1×10^5^ cells per well, treated with or without 5 µg/ml IFNAR1-blocking antibody (Leinco Technologies, Clone MAR1-5A3, I-401) to report IFN-I levels, and incubated overnight with 50 μl of neutrophil or alveolar macrophage supernatant. Released luciferase activity was detected using a QUANTI-Luc™ 4 Lucia/Gaussia Luciferase Activity Kit (InvivoGen) as per the manufacturer’s instructions, and luminescence was read on a Synergy HT plate reader (BioTek).

### Immunofluorescence microscopy

4% paraformaldehyde (PFA) fixed coverslips were washed with PBS and permeabilized in PBS with 2% FBS and 0.1% saponin for 30 minutes. To detect histone citrullination (H3Cit) in neutrophil extracellular traps (NETs), coverslips were stained with a rabbit anti-histone H3 citrulline R17 antibody (Abcam, ab219407, 2 µg/ml); to detect IFNα, coverslips were stained with a rabbit anti-IFNα polyclonal antibody (Invitrogen, PA5-119649, 1:200); to detect fibrillarin, coverslips were stained with chicken anti-fibrillarin (Invitrogen, PA5-143565, 1:500); to detect DDX21, coverslips were stained with rabbit anti-DDX21 (Novus Biologicals, NB100-1718, 2 µg/ml). Following incubation with primary antibody at 4°C overnight, coverslips were washed in PBS with 2% FBS and 0.1% saponin, and incubated with the appropriate secondary antibodies: donkey anti-rabbit Alexa Fluor 555 (Invitrogen) and/or goat anti-chicken Alexa Fluor 647 (Invitrogen), at 1:200 for 1 hour at room temperature. DNA was labeled with 5 μg/ml Hoechst 33342 (Invitrogen). Confocal images were acquired on a Nikon AXR confocal microscope coupled with NIS software at 20x magnification for wide field, or under 60x oil immersion for z-stack confocal images. Image quantification was performed using ImageJ FIJI, where the DNA channel in each slice of a z-stack was subjected to thresholding to generate a mask of the nucleus. IFNα staining in the nucleus was quantified by reporting the raw integrated intensity of IFNα signal within the nuclear mask, normalized to the area of the nuclear mask. Nucleolar size was quantified by reporting fibrillarin staining area within the nuclear mask, normalized to the area of the nuclear mask.

### HoxB8 progenitor cell line production and culture

The HoxB8 granulocyte-monocyte progenitor cell line was generated by expression of an estrogen receptor-homeobox B8 (ER-Hoxb8) gene fusion as previously described^33^. In brief, bone marrow cells from a C57BL/6J mouse were lineage depleted (Lineage Cell Depletion Kit, Miltenyi) and retroviral transduced with an MSCVneo-HA-ER-Hoxb8 construct via spinoculation at 1500□×□g for 1.5 h at 37°C, in the presence of 1:1000 Lipofectamine 2000 (Invitrogen). Transduced HoxB8 progenitor cells were maintained in 5% CO_2_ at 37°C, in HoxB8 progenitor media (RPMI 1640 media containing 10% FBS, 10ng/ml G-CSF (MedChem Express), 1□μM β-estradiol (Sigma), and 10 ng/ml stem cell factor (SCF) from conditioned supernatant of SCF-producing Chinese hamster ovary (CHO) cells kindly provided by Dr. David Sykes (Massachusetts General Hospital, MA)). Removal of β-estradiol from the cell media abolishes nuclear translocation of the ER-HoxB8 fusion protein and results in differentiation of the progenitor cells into mature HoxB8 neutrophils, and by 4 days after estradiol removal, differentiated HoxB8 neutrophils were used for experiments and infection as described.

### Nucleolar proximity labeling by APEX2

GFP-APEX2-NIK3x-expressing HoxB8 progenitor cells were generated by lentiviral transduction as previously described. In brief, HEK293T cells were seeded in 6-well plates the day before transfection and transfected at 80-90% confluency using 15 μl of Lipofectamine 2000, 250 μl Opti-MEM, and a DNA mixture of 0.5 μg of pMD2.G (Addgene #12259), 1 μg of psPAX2 (Addgene #12260), and 1.5 μg of GFP-APEX2-NIK3x plasmid (modified from Addgene #129274 to replace the CMV promoter with an SFFV promoter by Gibson assembly). Cells were incubated at 37°C with 5% CO_2_ for 4 hours, and then fresh HoxB8 progenitor media was added to dilute out lipofectamine. Lentivirus was harvested 2 days after the start of transfection and filtered through a 0.45 µm mixed cellulose esters (MCE) membrane (Millipore Sigma). Lentiviral spinfection was performed by centrifuging lentivirus-containing supernatant with HoxB8 progenitor cells in 12-well plates at 1,000×g for 2 hours at 37°C, in HoxB8 progenitor media and 10 μg/ml polybrene. Spinfected cells were incubated at 37°C with 5% CO_2_ for 4 hours, and then fresh HoxB8 progenitor media was added to dilute out polybrene. 3 days after lentiviral transduction, cell culture medium was replaced with HoxB8 progenitor media containing 2 μg/ml of blasticidin for selection of GFP-APEX2-NIK3x-expressing cells.

For proximity labeling, 500 μM biotin tyramide (MedChem Express) was spiked in cultures of GFP-APEX2-NIK3x-expressing HoxB8 progenitors or neutrophils, to a final concentration of 500 μM. After 60 minutes of incubation, 1×10^7^ cells were collected in conical tubes, and 100 mM H_2_O_2_ in PBS was added to a final concentration of 1 mM to each tube. After 1 minute of gentle agitation, the reaction was rapidly quenched with ice-cold quencher solution (10 mM sodium ascorbate (Sigma), 5 mM Trolox (MedChem Express), 10 mM sodium azide in PBS). Cells were spun down and washed with ice-cold quencher solution before pelleting and lysis by adding 500 µL of lysis buffer (1% SDS, 50 mM Tris-HCl pH 8.0, 10 mM sodium ascorbate, 5 mM trolox, 1 mM PMSF (GoldBio), 1:1000 benzonase (Sigma, E1014), Pierce EDTA-free Protease Inhibitor (ThermoFisher, A32955), 1:100 HALT protease inhibitor cocktail (ThermoFisher, 78441), and phosphatase inhibitor cocktails (Sigma, P5726, P0044)). Lysates were clarified by centrifugation at 10,000xg for 10 minutes at 4°C. Clarified supernatants were transferred to fresh low-binding tubes prior to streptavidin bead-based biotin enrichment. 100 μl of streptavidin beads (ThermoFisher, 65601) were washed 2 times in lysis buffer, added to each sample, and incubated with rotation for 2 hours for protein binding. Protein-bound beads were then washed with ice-cold buffers as follows: 2 times with lysis buffer, 1 time with 1M KCl, 1 time with 0.1 M Na_2_CO_3_, 1 time with 2 M Urea in 10 mM Tris-HCl (pH 8), and 1 time with PBS.

### Western blotting

Neutrophils were lysed in 1× RIPA buffer (Cell Signaling Technology, 9806) containing Pierce EDTA-free Protease Inhibitor, 1:100 HALT protease inhibitor cocktail, 1□mM PMSF, and phosphatase inhibitor cocktails. Lysates were diluted with 4× Laemmli buffer and immediately boiled for 20□minutes. Protein-bound streptavidin beads were directly added to 1x Laemmli buffer and boiled for 20 minutes. Protein samples were resolved on a 4–12% NuPAGE Bis-Tris gel (ThermoFisher), transferred to PVDF membrane (Millipore Sigma), blocked with 5% milk, and cut at 50 kDa. The <50kDa portion of the membrane was blotted with anti-IFNα (Invitrogen, PA5-119649, 1:1000), and the >50kDa portion was blotted with anti-HSP60 (Novus Biologicals, NBP1-77397SS, 1:1000), followed by goat anti-rabbit-horseradish peroxidase (HRP) (Thermo Fisher, 31460) incubation. HRP was detected with SuperSignal West Dura ECL substrate (ThermoFisher, 34075) and imaged on a ChemiDoc System (BioRad).

### RNA isolation and sequencing

RNA samples were collected using Trizol Reagent (Invitrogen), followed by extraction using chloroform. RNA was isolated using Direct-zol RNA Miniprep Plus kit (Zymo Research) as per the manufacturer’s instructions. For RNAseq, library preparation was performed at Washington University Genomic Technology Access Center (GTAC). In brief, RNA was fragmented in reverse transcriptase buffer while ribosomal RNA was blocked using FastSelect reagents (Qiagen), and mRNA was reverse transcribed to yield cDNA using SuperScript III RT enzyme (Life Technologies), indexed, pooled, and sequenced on an Illumina NovaSeq X Plus (Illumina, USA) using paired end reads extending 150 bases. Basecalls and demultiplexing were performed with Illumina’s bcl2fastq software with a maximum of one mismatch in the indexing read. RNA-seq reads were then aligned to the Ensembl release 101 primary assembly with STAR version 2.7.9a1^63^. Gene counts were derived from the number of uniquely aligned unambiguous reads by Subread:featureCount version 2.0.32^64^. Isoform expression of known Ensembl transcripts was quantified with Salmon version 1.5.23. Sequencing performance was assessed for the total number of aligned reads, the total number of uniquely aligned reads, and features detected. The ribosomal fraction, known junction saturation, and read distribution over known gene models were quantified with RSeQC version 4.04^65^. Differential expression analysis was performed in R version 4.5.1 using DESeq2 BioC 3.22 Version 1.50.2^66^ to analyze differences between conditions, and the results were filtered for only those genes with Benjamini-Hochberg false-discovery rate adjusted p-values less than or equal to 0.05. The volcano plot was generated in R by plotting the −log_10_(P-value) against the log_2_FC.

### Pre-rRNA detection by qRT-PCR

For qRT-PCR, cDNA was synthesized using the SuperScript III First-Strand kit (Invitrogen) using oligo(dT), and used for quantitative PCR in a Real-Time PCR System (Bio-Rad CFX96). Target gene cDNAs was detected using the iTaq Universal SYBR Green master mix (BioRad) and the following primer pairs: *Mouse Gapdh* forward CTTTGTCAAGCTCATTTCCTGG and reverse TCTTGCTCAGTGTCCTTGC, human *GAPDH* forward TGTCAGTGGTGGACCTGACCT and reverse AGGGGAGATTCAGTGTGGTG, mouse 45S pre-rRNA forward CTCTTAGATCGATGTGGTGCTC and reverse GCCCGCTGGCAGAACGAGAAG^67^, and human 47S/45S pre-rRNA 5’ external transcribed spacer (ETS) forward GGCGGTTTGAGTGAGACGAGA and reverse ACGTGCGCTCACCGAGAGCAG^47^. The 2^−ddCt^ method was used to determine gene expression fold change after normalization with the respective housekeeping genes.

### Flow cytometry

Flow cytometry was performed to assess HoxB8 neutrophil maturation, myeloperoxidase (MPO) content, and IFNα expression. Neutrophils were surface-stained at a 1:200 dilution with anti-mouse CD16/32 Fc blocker (BioLegend), GR-1-Pacific Blue (BioLegend, 108429), and CD11b-Brilliant Violet 650 (BioLegend, 101239) in PBS with 2% FBS. Surface-stained neutrophils were fixed in 4% PFA and permeabilized in FOXP3 perm buffer (BioLegend) containing 1:200 Fc blocker and 5% normal goat serum (Sigma). Permeabilized cells were stained with MPO-Alexa Fluor 488 (Abcam, ab322168, 1:200) and rabbit anti-IFNα (Invitrogen, PA5-119649, 1:200), followed by a goat anti-rabbit Alexa Fluor 700 secondary (Invitrogen, 1:800). Flow cytometry was performed on an Aurora spectral flow cytometer (Cytek Biosciences, 4 laser 16V-14B-10YG-8R configuration), spectral unmixing was performed using SpectroFlo (Cytek Biosciences), and data were analyzed using FlowJo software (BD Biosciences).

### Imaging flow cytometry

To visualize IFNα localization in HoxB8 neutrophils, 4% PFA fixed cells were permeabilized and stained with anti-IFNα (Invitrogen, PA5-119649) and anti-fibrillarin (Invitrogen, PA5-143565), followed by goat anti-rabbit Alexa Fluor 488 and goat anti-chicken Alexa Fluor 647 secondaries. DNA was labeled with 5 μg/ml Hoechst 33342. To visualize IFNα localization in heat-killed Mtb-treated mouse neutrophils, cells were surface-stained with Ly6G-Pacific Blue (BioLegend, 127612), fixed in 4% PFA, and permeabilized for staining with anti-IFNα, followed by a by goat anti-rabbit Alexa Fluor 647 secondary (Invitrogen). DNA was labeled with 0.1µM Sytox green (Invitrogen). Imaging flow cytometry was performed on an ImageStreamX Mark II instrument operated by INSPIRE software (Cytek Biosciences). A total of 5000-10,000 singlet cell events were collected for each sample, and single-stained controls were collected with the same settings for use in compensation.

Data analysis was performed using the IDEAS Application v6.2 software (Cytek Biosciences). For IFNα localization analysis, in focus and single cell events were selected, and cell compartment masks were created as described in Supplemental Figures 2C and 3A. IFNα bright detail intensity was determined within each mask to report the percentage of IFNα signal in each compartment. Machine learning to identify features correlating with nucleolar IFNα expression was conducted in IDEAS Application v6.4 software (Cytek Biosciences). Nucleolar IFNα-high and IFNα-low cells were gated on separately, and the Hoechst and brightfield channels were selected for correlation machine learning analysis. The top features correlating to nucleolar IFNα expression were “Intensity_Valley(Morphology(Hoechest_Mask,Hoechest),Hoechest,3)_Hoechst”, which we denoted as nuclear segmentation, when using all features for analysis; and “Length_Object(M01,Hoechst,Tight)”, which we denoted as nuclear elongation, when restricting to shape and size features for analysis.

### Statistical analysis

All data are from at least two independent experiments, and samples represent replicates of individual wells of cells. Statistical analyses were performed with Prism (v10.6.0; GraphPad Software), using unpaired two-tailed Student’s t tests to compare between two conditions, one-way ANOVA with Tukey’s correction to perform multiple comparisons, or with two-way ANOVA with Tukey’s correction to perform multiple comparisons in time course experiments with two conditions. When used, center values and error bars represent mean ± SEM, and p < 0.05 was considered significant. In all figures, significant differences are indicated by asterisks: * p<0.05, ** p<0.01, *** p<0.001, **** p<0.0001. Only significant differences are noted in the figures unless otherwise specified, where nonsignificant differences were denoted as ns.

## Notes

### Competing Interest Statement

The authors have declared no competing interest.

### Summary of Updates

Title and abstract updated to better encapsulate scope of the study.

## References

1 McNab, F., Mayer-Barber, K., Sher, A., Wack, A. & O’Garra, A. Type I interferons in infectious disease. Nat Rev Immunol 15, 87–103 (2015). 10.1038/nri3787

2 Watson, R. O. et al. The Cytosolic Sensor cGAS Detects Mycobacterium tuberculosis DNA to Induce Type I Interferons and Activate Autophagy. Cell Host Microbe 17, 811–819 (2015). 10.1016/j.chom.2015.05.004

3 Gibbert, K., Schlaak, J. F., Yang, D. & Dittmer, U. IFN-alpha subtypes: distinct biological activities in anti-viral therapy. Br J Pharmacol 168, 1048–1058 (2013). 10.1111/bph.12010

4 van Pesch, V., Lanaya, H., Renauld, J. C. & Michiels, T. Characterization of the murine alpha interferon gene family. J Virol 78, 8219–8228 (2004). 10.1128/JVI.78.15.8219-8228.2004

5 Hardy, M. P., Owczarek, C. M., Jermiin, L. S., Ejdeback, M. & Hertzog, P. J. Characterization of the type I interferon locus and identification of novel genes. Genomics 84, 331–345 (2004). 10.1016/j.ygeno.2004.03.003

6 Moll, H. P., Maier, T., Zommer, A., Lavoie, T. & Brostjan, C. The differential activity of interferon-alpha subtypes is consistent among distinct target genes and cell types. Cytokine 53, 52–59 (2011). 10.1016/j.cyto.2010.09.006

7 Londe, A. C., Fernandez-Ruiz, R., Julio, P. R., Appenzeller, S. & Niewold, T. B. Type I Interferons in Autoimmunity: Implications in Clinical Phenotypes and Treatment Response. J Rheumatol 50, 1103–1113 (2023). 10.3899/jrheum.2022-0827

8 Berry, M. P. et al. An interferon-inducible neutrophil-driven blood transcriptional signature in human tuberculosis. Nature 466, 973–977 (2010). 10.1038/nature09247

9 Nair, S. et al. Irg1 expression in myeloid cells prevents immunopathology during M. tuberculosis infection. J Exp Med 215, 1035–1045 (2018). 10.1084/jem.20180118

10 Munson, C. W. & Kaushal, D. Type I interferons in tuberculosis pathogenesis and prevention. Trends Microbiol 34, 293–304 (2026). 10.1016/j.tim.2025.11.003

11 Niazi, M. K. et al. Lung necrosis and neutrophils reflect common pathways of susceptibility to Mycobacterium tuberculosis in genetically diverse, immune-competent mice. Dis Model Mech 8, 1141–1153 (2015). 10.1242/dmm.020867

12 Esaulova, E. et al. The immune landscape in tuberculosis reveals populations linked to disease and latency. Cell Host Microbe 29, 165–178 e168 (2021). 10.1016/j.chom.2020.11.013

13 Moreira-Teixeira, L. et al. Type I IFN exacerbates disease in tuberculosis-susceptible mice by inducing neutrophil-mediated lung inflammation and NETosis. Nat Commun 11, 5566 (2020). 10.1038/s41467-020-19412-6

14 Chowdhury, C. S. et al. Type I IFN-mediated NET release promotes Mycobacterium tuberculosis replication and is associated with granuloma caseation. Cell Host Microbe 32, 2092–2111 e2097 (2024). 10.1016/j.chom.2024.11.008

15 Kinsella, R. L. et al. ATG5 suppresses type I IFN-dependent neutrophil effector functions during Mycobacterium tuberculosis infection in mice. Nat Microbiol 10, 1323–1339 (2025). 10.1038/s41564-025-01988-8

16 Lyadova, I. Type I interferons in bacterial diseases: myeloid cells at the crossroads of protection and pathology. Front Immunol 16, 1717370 (2025). 10.3389/fimmu.2025.1717370

17 Pylaeva, E. et al. Detrimental Effect of Type I IFNs During Acute Lung Infection With Pseudomonas aeruginosa Is Mediated Through the Stimulation of Neutrophil NETosis. Front Immunol 10, 2190 (2019). 10.3389/fimmu.2019.02190

18 Branchett, W. J. et al. Type I IFN drives neutrophil swarming, impeding lung T cell-macrophage interactions and TB control. J Exp Med 222 (2025). 10.1084/jem.20250466

19 Lee, A. M., Laurent, P., Nathan, C. F. & Barrat, F. J. Neutrophil-plasmacytoid dendritic cell interaction leads to production of type I IFN in response to Mycobacterium tuberculosis. Eur J Immunol 54, e2350666 (2024). 10.1002/eji.202350666

20 Kotov, D. I. et al. Early cellular mechanisms of type I interferon-driven susceptibility to tuberculosis. Cell 186, 5536–5553 e5522 (2023). 10.1016/j.cell.2023.11.002

21 TenBarge, E. G. et al. A TLR7/9-IFNalpha-LDHB axis drives vital NET release and compromises antibacterial defense in lupus. bioRxiv (2025). 10.1101/2025.10.07.680928

22 Tamassia, N. et al. Activation of an immunoregulatory and antiviral gene expression program in poly(I:C)-transfected human neutrophils. J Immunol 181, 6563–6573 (2008). 10.4049/jimmunol.181.9.6563

23 Lindau, D. et al. TLR9 independent interferon alpha production by neutrophils on NETosis in response to circulating chromatin, a key lupus autoantigen. Ann Rheum Dis 73, 2199–2207 (2014). 10.1136/annrheumdis-2012-203041

24 Yang, E. & Li, M. M. H. All About the RNA: Interferon-Stimulated Genes That Interfere With Viral RNA Processes. Front Immunol 11, 605024 (2020). 10.3389/fimmu.2020.605024

25 Matta, S. K. et al. Toxoplasma gondii effector TgIST blocks type I interferon signaling to promote infection. Proc Natl Acad Sci U S A 116, 17480–17491 (2019). 10.1073/pnas.1904637116

26 Kendall, R. L., Ray, J. L., Hamilton, R. F., Jr. & Holian, A. Self-replicating murine ex vivo cultured alveolar macrophages as a model for toxicological studies of particle-induced inflammation. Toxicol Appl Pharmacol 461, 116400 (2023). 10.1016/j.taap.2023.116400

27 Wassermann, R. et al. Mycobacterium tuberculosis Differentially Activates cGAS- and Inflammasome-Dependent Intracellular Immune Responses through ESX-1. Cell Host Microbe 17, 799–810 (2015). 10.1016/j.chom.2015.05.003

28 Cheng, Y. & Schorey, J. S. Mycobacterium tuberculosis-induced IFN-beta production requires cytosolic DNA and RNA sensing pathways. J Exp Med 215, 2919–2935 (2018). 10.1084/jem.20180508

29 Collins, A. C. et al. Cyclic GMP-AMP Synthase Is an Innate Immune DNA Sensor for Mycobacterium tuberculosis. Cell Host Microbe 17, 820–828 (2015). 10.1016/j.chom.2015.05.005

30 Shannon, J. G. & Hinnebusch, B. J. Characterization and CRISPR/Cas9-mediated genetic manipulation of neutrophils derived from Hoxb8-ER-immortalized myeloid progenitors. J Leukoc Biol 114, 42–52 (2023). 10.1093/jleuko/qiad036

31 Lacy, P. Mechanisms of degranulation in neutrophils. Allergy Asthma Clin Immunol 2, 98–108 (2006). 10.1186/1710-1492-2-3-98

32 van Grinsven, E., Mukherjee, A. K. & Udalova, I. A. Shaping neutrophil morphology and function: the importance of a segmented nucleus. Nat Rev Immunol 26, 269–283 (2026). 10.1038/s41577-025-01237-3

33 Wang, G. G. et al. Quantitative production of macrophages or neutrophils ex vivo using conditional Hoxb8. Nat Methods 3, 287–293 (2006). 10.1038/nmeth865

34 Fazal, F. M. et al. Atlas of Subcellular RNA Localization Revealed by APEX-Seq. Cell 178, 473–490 e426 (2019). 10.1016/j.cell.2019.05.027

35 Pan, Y. H. et al. Pre-rRNA spatial distribution and functional organization of the nucleolus. Nature 646, 227–235 (2025). 10.1038/s41586-025-09412-1

36 Quinodoz, S. A. et al. Mapping and engineering RNA-driven architecture of the multiphase nucleolus. Nature 644, 557–566 (2025). 10.1038/s41586-025-09207-4

37 Riback, J. A. et al. Viscoelasticity and advective flow of RNA underlies nucleolar form and function. Mol Cell 83, 3095–3107 e3099 (2023). 10.1016/j.molcel.2023.08.006

38 Shen, C. et al. Nuclear segmentation facilitates neutrophil migration. J Cell Sci 136 (2023). 10.1242/jcs.260768

39 Szaflarski, W. et al. Early rRNA processing is a stress-dependent regulatory event whose inhibition maintains nucleolar integrity. Nucleic Acids Res 50, 1033–1051 (2022). 10.1093/nar/gkab1231

40 Detres, D., Camacho-Badillo, A. & Calo, E. A pH-Centric Model of Nucleolar Activity and Regulation. J Mol Biol 437, 169136 (2025). 10.1016/j.jmb.2025.169136

41 Wang, M., Bokros, M., Theodoridis, P. R. & Lee, S. Nucleolar Sequestration: Remodeling Nucleoli Into Amyloid Bodies. Front Genet 10, 1179 (2019). 10.3389/fgene.2019.01179

42 Hua, L., Yan, D., Wan, C. & Hu, B. Nucleolus and Nucleolar Stress: From Cell Fate Decision to Disease Development. Cells 11 (2022). 10.3390/cells11193017

43 Zhu, Y. et al. Comprehensive characterization of neutrophil genome topology. Genes Dev 31, 141–153 (2017). 10.1101/gad.293910.116

44 Burn, G. L., Foti, A., Marsman, G., Patel, D. F. & Zychlinsky, A. The Neutrophil. Immunity 54, 1377–1391 (2021). 10.1016/j.immuni.2021.06.006

45 Pandey, S., Kawai, T. & Akira, S. Microbial sensing by Toll-like receptors and intracellular nucleic acid sensors. Cold Spring Harb Perspect Biol 7, a016246 (2014). 10.1101/cshperspect.a016246

46 Cervantes, J. L. MyD88 in Mycobacterium tuberculosis infection. Med Microbiol Immunol 206, 187–193 (2017). 10.1007/s00430-017-0495-0

47 Aryan, F. et al. Nucleolus activity-dependent recruitment and biomolecular condensation by pH sensing. Mol Cell 83, 4413–4423 e4410 (2023). 10.1016/j.molcel.2023.10.031

48 Yang, K., Yang, J. & Yi, J. Nucleolar Stress: hallmarks, sensing mechanism and diseases. Cell Stress 2, 125–140 (2018). 10.15698/cst2018.06.139

49 Nalabothula, N., Indig, F. E. & Carrier, F. The Nucleolus Takes Control of Protein Trafficking Under Cellular Stress. Mol Cell Pharmacol 2, 203–212 (2010).

50 Potapova, T. A. et al. Distinct states of nucleolar stress induced by anticancer drugs. Elife 12 (2023). 10.7554/eLife.88799

51 Yang, Y. M. & Karbstein, K. Ribosome Assembly and Repair. Annu Rev Cell Dev Biol 40, 241–264 (2024). 10.1146/annurev-cellbio-111822-113326

52 Shafqat, A. et al. Neutrophil extracellular traps and long COVID. Front Immunol 14, 1254310 (2023). 10.3389/fimmu.2023.1254310

53 Cross, A. L. et al. Circulating neutrophil extracellular trap-forming neutrophils in rheumatoid arthritis exacerbation are majority dual endothelin-1/signal peptide receptor+ subtype. Clin Exp Immunol 218, 163–168 (2024). 10.1093/cei/uxae072

54 Vance, R. E. Tuberculosis as an unconventional interferonopathy. Curr Opin Immunol 92, 102508 (2025). 10.1016/j.coi.2024.102508

55 Xu, S. et al. Pore-forming activity of S. pneumoniae pneumolysin disrupts the paracellular localization of the epithelial adherens junction protein E-cadherin. Infect Immun 91, e0021323 (2023). 10.1128/iai.00213-23

56 Greene, N. G., Narciso, A. R., Filipe, S. R. & Camilli, A. Peptidoglycan Branched Stem Peptides Contribute to Streptococcus pneumoniae Virulence by Inhibiting Pneumolysin Release. PLoS Pathog 11, e1004996 (2015). 10.1371/journal.ppat.1004996

57 Shields-Cutler, R. R. et al. Human Urinary Composition Controls Antibacterial Activity of Siderocalin. J Biol Chem 290, 15949–15960 (2015). 10.1074/jbc.M115.645812

58 Chen, S. L. et al. Identification of genes subject to positive selection in uropathogenic strains of Escherichia coli: a comparative genomics approach. Proc Natl Acad Sci U S A 103, 5977–5982 (2006). 10.1073/pnas.0600938103

59 Steed, A. L. et al. The microbial metabolite desaminotyrosine protects from influenza through type I interferon. Science 357, 498–502 (2017). 10.1126/science.aam5336

60 Szretter, K. J. et al. The innate immune adaptor molecule MyD88 restricts West Nile virus replication and spread in neurons of the central nervous system. J Virol 84, 12125–12138 (2010). 10.1128/JVI.01026-10

61 Adachi, O. et al. Targeted disruption of the MyD88 gene results in loss of IL-1- and IL-18-mediated function. Immunity 9, 143–150 (1998). 10.1016/s1074-7613(00)80596-8

62 Sato, M. et al. Distinct and essential roles of transcription factors IRF-3 and IRF-7 in response to viruses for IFN-alpha/beta gene induction. Immunity 13, 539–548 (2000). 10.1016/s1074-7613(00)00053-4

63 Dobin, A. et al. STAR: ultrafast universal RNA-seq aligner. Bioinformatics 29, 15–21 (2013). 10.1093/bioinformatics/bts635

64 Liao, Y., Smyth, G. K. & Shi, W. featureCounts: an efficient general purpose program for assigning sequence reads to genomic features. Bioinformatics 30, 923–930 (2014). 10.1093/bioinformatics/btt656

65 Wang, L., Wang, S. & Li, W. RSeQC: quality control of RNA-seq experiments. Bioinformatics 28, 2184–2185 (2012). 10.1093/bioinformatics/bts356

66 Love, M. I., Anders, S., Kim, V. & Huber, W. RNA-Seq workflow: gene-level exploratory analysis and differential expression. F1000Res 4, 1070 (2015). 10.12688/f1000research.7035.1

67 Watada, E. et al. Age-Dependent Ribosomal DNA Variations in Mice. Mol Cell Biol 40 (2020). 10.1128/MCB.00368-20

