## Supplemental Figures for "Neutrophils repurpose the nucleolus as a cytokine reservoir and secretory organelle"

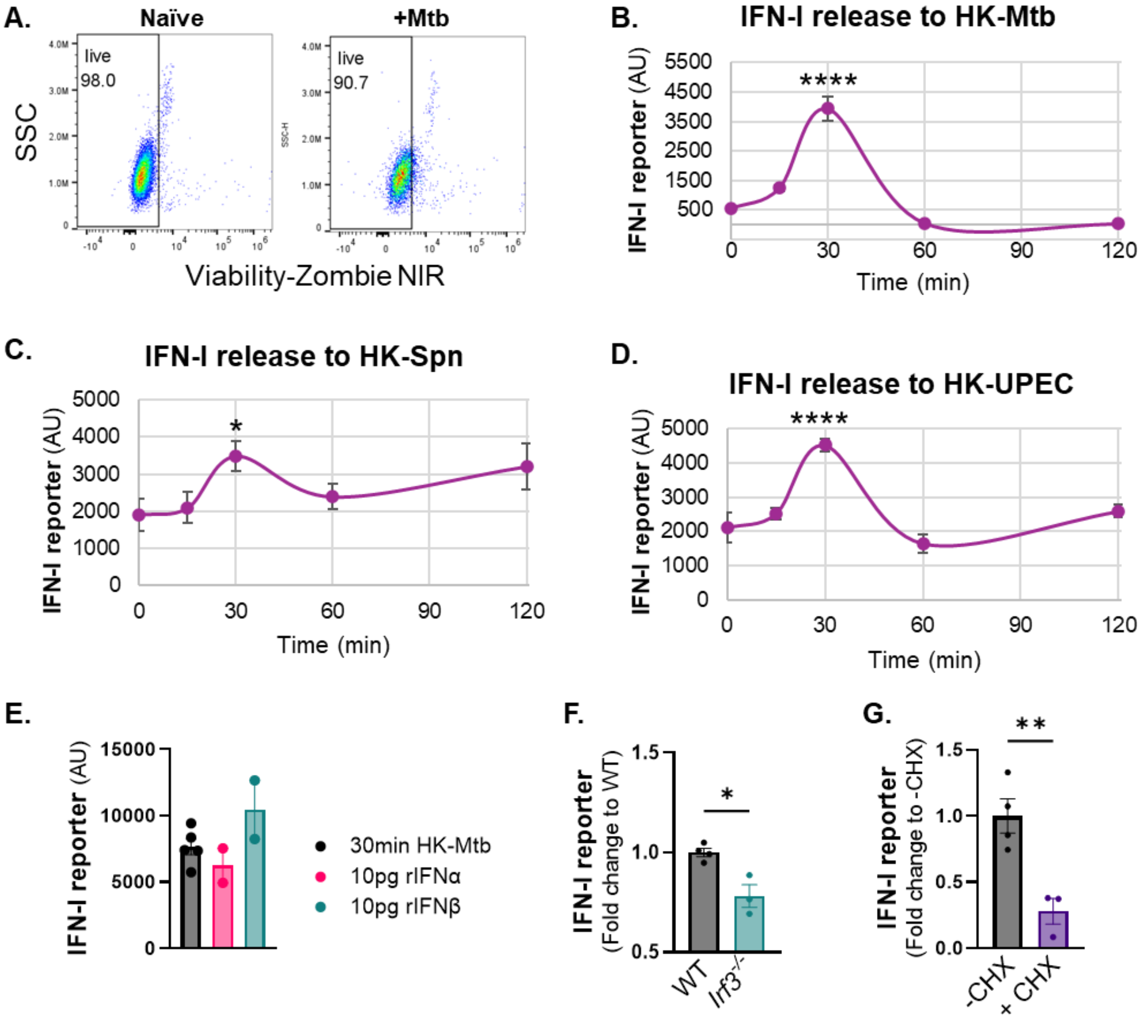


**Supplemental Figure 1. Heat-killed bacteria are sufficient to trigger rapid IFN-I release from neutrophils. A.** FACS analysis of viability for human neutrophils naïve or 120 mpi with Mtb at an MOI of 5. **B-D.** Total IFNAR-dependent luciferase activity from ISRE reporter cells exposed to supernatants collected from mouse neutrophils treated with heat-killed (HK) Mtb (A), Spn (B), or UPEC (C). **E.** Total IFNAR-dependent luciferase activity from ISRE reporter cells exposed to supernatants collected from mouse neutrophils treated with heat-killed Mtb for 30 min, or directly to 10pg of recombinant IFNα or IFNβ. **F.** Total IFNAR-dependent luciferase activity from ISRE reporter cells exposed to supernatants collected from WT or *Irf3*$^{-/-}$ mouse neutrophils at 120 minutes post exposure to poly(I:C). **G.** Total IFNAR-dependent luciferase activity from ISRE reporter cells exposed to supernatants collected from human neutrophils at 120 minutes post exposure to poly(I:C), with 10 µg/mL cycloheximide pre-treatment as indicated. Each data point is the average of independent wells of cells (A-C), or represents independent wells of cells (D-F) in a representative experiment from at least two independent experiments. Error bars represent mean ± SEM. Statistical differences to zero timepoint, between genotypes, or between treatments were determined using ordinary one-way ANOVA (A-C) with Tukey's post-hoc test, or Student's t-test (E-F): *p < 0.05, **p < 0.01, ****p < 0.0001.

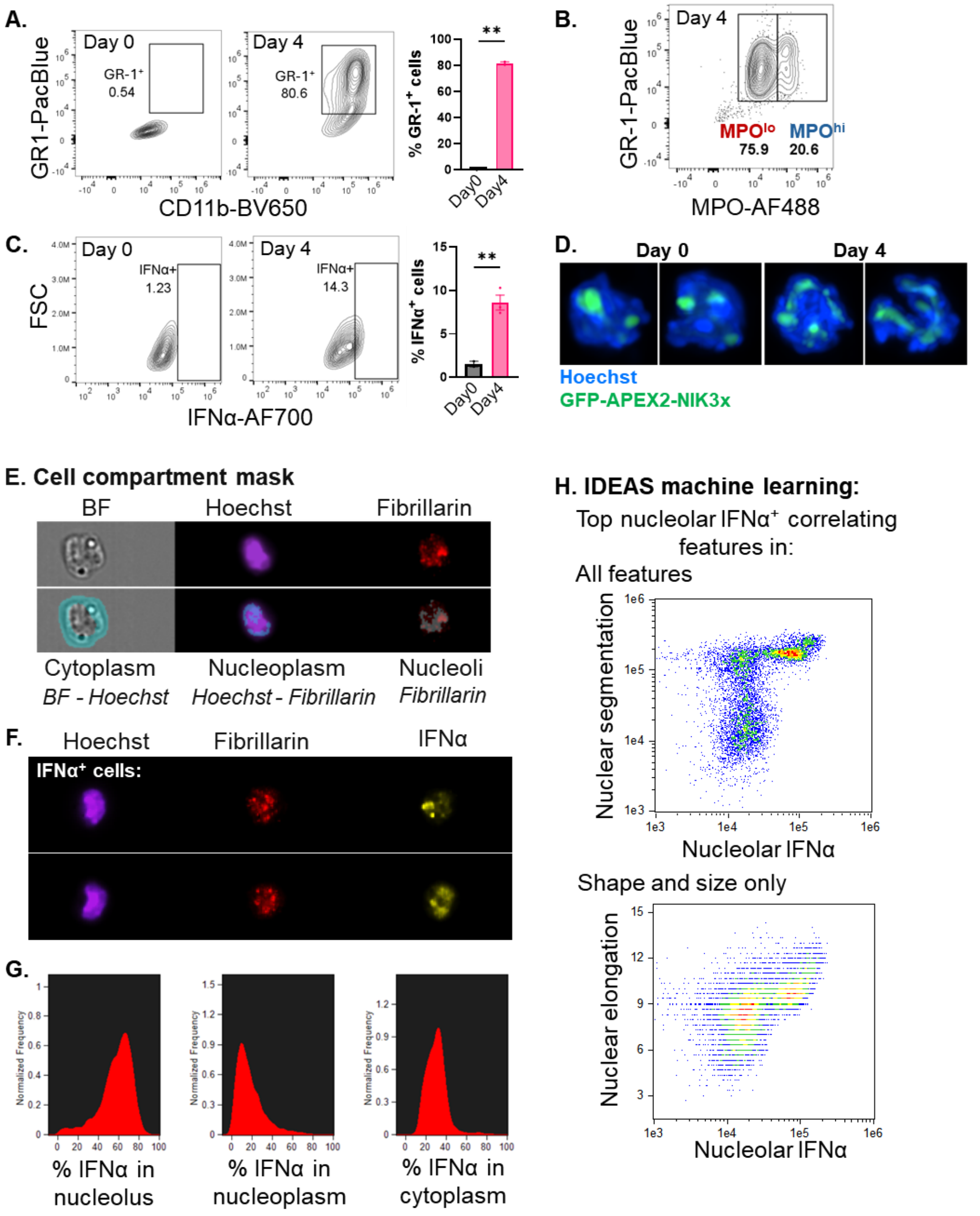


**Supplemental Figure 2. IFNα is stored in the nucleoli of mature neutrophils. A-C.** FACS analysis of CD11b and GR-1 expression (A), intracellular MPO expression (B), or intracellular IFNα expression (C) in HoxB8 progenitors (Day 0) or HoxB8 neutrophils (Day 4). **D.** Immunofluorescence confocal microscopy of GFP-APEX2-NIK3x-expressing HoxB8 progenitors (Day 0) and HoxB8 neutrophils (Day 4). Images show DNA (Hoechst) and GFP-tagged APEX2-NIK3x. **E-H.** Imagestream analysis of naïve Hoxb8 neutrophils. Images show brightfield, DNA (Hoechst), fibrillarin, and IFNα, with masking scheme used to identify nucleoli, nucleoplasm, and cytoplasm (E), representative images of IFNα$^{+}$ cells (F), quantitation of percent IFNα signal in respective masks (G), and correlation of nuclear segmentation and elongation with IFNα nucleolar expression as features identified by IDEAS machine learning (H). Each data point represents independent wells of cells in a representative experiment from at least two independent experiments. Error bars represent mean ± SEM. Statistical differences between Day 0 and Day 4 HoxB8 cells were determined using Student's t-test: $**p < 0.01$.

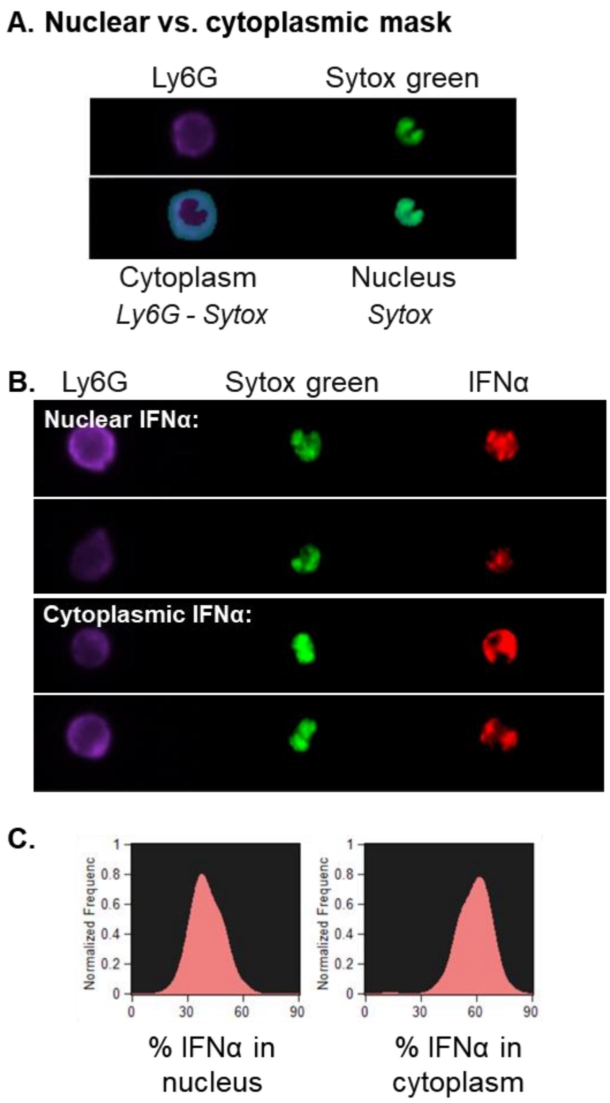


**Supplemental Figure 3. Neutrophil IFNα is released into the cytoplasm before extracellular secretion. A-C** Imagestream analysis of mouse neutrophils treated with heat-killed Mtb. Images show Ly6G, DNA (sytox green), and IFNα, with masking scheme used to identify nucleus and cytoplasm (A), representative images of cells with nuclear or cytoplasmic IFNα (B), and quantitation of percent IFNα signal in respective masks (C). Shown is representative of at least two independent experiments.

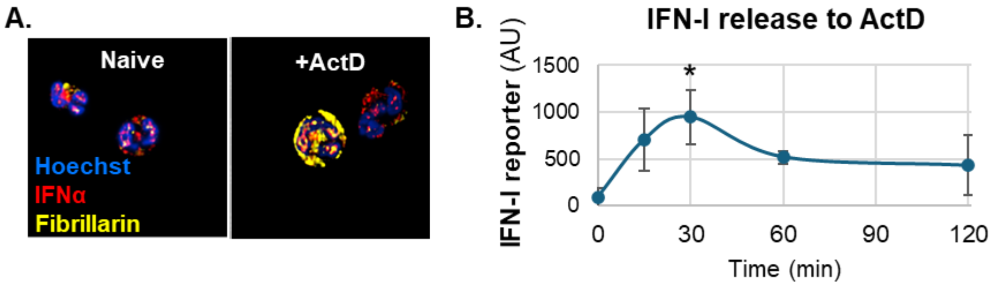


**Supplemental Figure 4. Actinomycin D is sufficient to trigger IFN-I release from human neutrophils. A.** Immunofluorescence confocal microscopy of human neutrophils naïve or treated with heat-killed Mtb. Images show DNA (Hoechst), IFNα, and fibrillarin. **B.** Total IFNAR-dependent luciferase activity from ISRE reporter cells exposed to supernatants collected from human neutrophils treated with 50nM actinomycin D. Each data point is the average of independent wells of infected cells in a representative experiment from at least two independent experiments. Error bars represent mean ± SEM. Statistical differences to the zero timepoint were determined using ordinary one-way ANOVA with Tukey's post-hoc test: $^*p < 0.05$.
