## Supplemental Table for "Neutrophils repurpose the nucleolus as a cytokine reservoir and secretory organelle"

**Supplemental table 1.** List of genes differentially expressed in Mtb-infected WT and *Ifnar^-/-^* mouse neutrophils.

| **Gene** | **Base Mean** | **log2FoldChange** | **lfcSE** | **Stat** | **P_adj._** |
| --- | --- | --- | --- | --- | --- |
| Hist1h4m | 177.051 | -3.659 | 0.368 | -9.956 | 2.36E-23 |
| 5830416I19Rik | 146.189 | 2.869 | 0.320 | 8.972 | 2.90E-19 |
| Cryzl1 | 1138.952 | 1.792 | 0.219 | 8.178 | 2.88E-16 |
| Gm10785 | 231.241 | 2.151 | 0.264 | 8.158 | 3.42E-16 |
| Epx | 78.414 | 3.964 | 0.508 | 7.801 | 6.14E-15 |
| H2-Q6 | 81.377 | -2.991 | 0.386 | -7.749 | 9.25E-15 |
| H2-Q7 | 114.256 | -2.313 | 0.304 | -7.601 | 2.94E-14 |
| Itsn1 | 3982.230 | 1.295 | 0.176 | 7.343 | 2.09E-13 |
| Itgb2l | 2943.934 | -1.261 | 0.202 | -6.237 | 4.45E-10 |
| Donson | 1002.311 | 1.347 | 0.215 | 6.253 | 4.03E-10 |
| Irf9 | 507.383 | -1.318 | 0.226 | -5.843 | 5.12E-09 |
| H2-Q5 | 70.402 | -2.176 | 0.373 | -5.835 | 5.39E-09 |
| Plxna1 | 454.426 | 1.170 | 0.207 | 5.637 | 1.73E-08 |
| Rell1 | 1336.396 | 1.048 | 0.187 | 5.610 | 2.03E-08 |
| Gm21451 | 31.309 | -2.744 | 0.511 | -5.370 | 7.87E-08 |
| Oasl2 | 643.841 | -1.490 | 0.278 | -5.355 | 8.53E-08 |
| Mpo | 44.443 | 2.380 | 0.446 | 5.342 | 9.20E-08 |
| Slfn8 | 87.643 | -1.986 | 0.386 | -5.138 | 2.78E-07 |
| Vmn2r96 | 123.615 | 1.891 | 0.369 | 5.126 | 2.97E-07 |
| H2bc7 | 77.834 | 1.545 | 0.337 | 4.578 | 4.69E-06 |
| Ighg2b | 156.423 | 1.579 | 0.347 | 4.545 | 5.48E-06 |
| Cbx5 | 139.933 | 1.392 | 0.313 | 4.445 | 8.79E-06 |
| Tmem181b-ps | 180.861 | 1.052 | 0.247 | 4.258 | 2.06E-05 |
| Irgm2 | 66.315 | -1.621 | 0.388 | -4.182 | 2.88E-05 |
| Trim30a | 126.355 | -1.406 | 0.339 | -4.148 | 3.35E-05 |
| Diaph3 | 54.868 | 1.675 | 0.405 | 4.133 | 3.59E-05 |
| Mmp13 | 40.184 | 1.718 | 0.434 | 3.955 | 7.67E-05 |
| E2f8 | 124.170 | 1.155 | 0.294 | 3.924 | 8.73E-05 |
| Jchain | 180.717 | 0.968 | 0.247 | 3.914 | 9.08E-05 |
| Abce1 | 87.832 | 1.398 | 0.360 | 3.878 | 1.05E-04 |
